# Integrative Single-Cell Analysis of Autism Spectrum Disorder Animal Models Reveal Convergent Transcriptomic Dysregulation Involved in Excitatory-Inhibitory Imbalance and Glial Disfunction

**DOI:** 10.1101/2025.05.05.651905

**Authors:** João V. Nani, Victor J. Duque, Alysson R. Muotri, André S. Mecawi

**Author notes:** Corresponding author: Prof. Dr. André S. Mecawi, *Ph.D.*, and Dr. João V. S. Nani, *Ph.D.

## Abstract

Autism Spectrum Disorder (ASD) presents profound clinical and etiological heterogeneity, complicating the identification of core pathophysiological mechanisms. Single-cell RNA sequencing (scRNA-seq) offers cellular resolution but integrating findings across diverse studies remains challenging. Here, we constructed a unified single-cell reference framework by integrating scRNA-seq data from 11 distinct genetic and environmental ASD animal models, encompassing over 300.000 cells across various brain regions and developmental stages. Comparative analyses revealed convergent differentially expressed genes (DEGs) across neuronal and glial populations. Cross-model comparisons validated the integration, showing significant concordance between the unified dataset and individual studies, particularly for neuronal populations, and demonstrating how environmental models like valproic acid exposure recapitulate some of the transcriptomic alterations seen in genetic models. Cell communication analyses support widespread excitatory-inhibitory imbalance and with predicted signaling involving ligands like *Pdgfa* and *Reln*. Furthermore, we identified significant glial dysfunction, notably downregulation of crucial functional genes in astrocytes and signatures of metabolic dysregulation in mature oligodendrocytes. Cross-referencing with the SFARI database confirmed significant overlap with high-confidence ASD risk genes, with notable dysregulated in specific cell types included *Ermn* (upregulated in multiple glia), *Foxg1* (downregulated in L5/6 NP neurons) and *Mef2c* (downregulated in MEIS2-like interneurons). Comparison with human scRNA-seq postmortem data revealed conserved dysregulation, highlighting enrichment of presynaptic/postsynaptic translation processes in neurons (implicating *CACNAIA*, *GRIN2B*, *CAMK2A*, ribosomal proteins) along with enrichment for neurodevelopmental disorder pathways in mature oligodendrocytes, involving *NRXN* and *DLGAP* gene networks. This integrative study provides unprecedented insight into the convergent cellular and molecular pathologies underlying ASD, establishing a valuable resource for understanding shared mechanisms and identifying new potential therapeutic targets.

## INTRODUCTION

Autism spectrum disorder (ASD) is a complex neurodevelopmental condition defined by a wide range of social, communicative, and behavioral challenges (Hirota et al., 2023). It is estimated that about 1 in 100 children around the globe receive an ASD diagnosis (Zeidan et al., 2022). The prevalence of ASD has risen in recent years, which may be partially explained by greater awareness and enhanced diagnostic criteria, although the multifaceted nature of ASD still poses challenges for diagnosis and intervention (Palinkas et al., 2019). The exact causes of ASD remain largely unknown, reflecting its multifactorial complexity and variability: different individuals show significant variations in symptom severity and treatment response (Hodges et al., 2020).

Genome-wide association studies (GWAS) have identified numerous genetic variants associated with ASD, including highly penetrant mutations in genes such as *CHD7*, *SHANK3*, and *FOXG1* (Casanova et al., 2016). However, most cases likely arise from complex interactions among multiple small-effect genes and environmental factors, including perinatal complications such as viral infections and hypoxia (Karimi et al., 2017). These factors contribute to the disorder’s heterogeneity and further complicates efforts to establish a uniform biological basis for ASD. Current ASD treatments, including behavioral therapy and pharmacological interventions that target specific symptoms, fall short of addressing the underlying biological mechanisms. This gap leaves a critical shortfall in our understanding and management of the disorder (Hirota et al., 2023) and underscores the importance of innovative approaches to studying ASD.

Recent advances in sequencing technologies, particularly single-cell RNA sequencing (scRNA-seq), have unlocked new opportunities for investigating transcriptomic changes in the brain associated with ASD. By enabling transcriptome analysis at the single-cell level, scRNA-seq reveals the intricate cellular heterogeneity of complex tissues like the brain and identifies specific cell subpopulations that may be dysregulated (Chehimi et al., 2023). This approach is especially relevant for ASD, where molecular alterations may occur in discrete subsets of neurons or glial cells that are often missed in bulk tissue analyses. Recent scRNA-seq studies have identified transcriptomic changes in certain cell types, such as excitatory neurons and activated microglia from the upper cortical layers, both of which appear crucial to ASD pathogenesis and correlate with clinical severity (Velmeshev et al., 2019). Altered synaptic signaling in projection neurons further indicates a direct connection between cortical circuits and ASD-related behaviors (Wamsley et al., 2024). Another study supports these findings by demonstrating a dysregulation of cortical circuits in individuals with ASD that spans multiple regions, including primary sensory areas like the visual cortex, progressing along an anterior-posterior gradient (Gandal et al., 2022).

Although using postmortem brain samples from individuals with ASD has yielded valuable insights, these methods come with significant limitations. ASD is a developmental condition, and postmortem analyses often reflect later life stages, which may not capture the critical changes occurring during neurodevelopment (Fetit et al., 2021). Furthermore, the subjects can be exposed to factors such as prolonged medication use, which can alter gene expression (LeClerc and Easley, 2015), as well as by neurodegenerative processes and other comorbidities, making it difficult to isolate changes solely associated with ASD (Kern et al., 2013). Animal models have long been a cornerstone of ASD research, providing valuable insights into the disorder’s genetic and environmental components and helping to address the limitations of human studies. These models are usually created through specific gene manipulations, such as knocking out or overexpressing genes implicated in ASD, or by using pharmacological methods to mimic environmental influences, and allow researchers to investigate specific brain regions, such as the cortex and hippocampus, and to examine a range of developmental stages from embryonic to adulthood (Sierra-Arregui et al., 2020).

Although these approaches have provided valuable insights, animal models studies are frequently limited to single experimental conditions or genetic modifications. Moreover, differences in methodologies and study focus complicate cross-study comparisons and data integration (Ryu et al., 2023). Recent advances in bioinformatics, particularly algorithms designed to correct for bench effect while preserving biologically relevant signals, now offer a unique opportunity to identify conserved differentially expressed genes (DEGs) across various paradigms or conditions (Hrovatin et al., 2023; Zhou et al., 2023), brain regions (Gandal et al., 2022), and developmental stages (Kim et al., 2020). Such integrative analyses would not only enable the discovery of shared molecular pathways central to ASD but also leverage the strengths of diverse experimental designs and conditions.

Therefore, a promising strategy to overcome these challenges is the integration of multiple scRNA-seq datasets from diverse ASD animal models. Here, we integrated single-cell RNA sequencing datasets from 11 distinct animal models representing diverse genetic and environmental factors relevant to ASD, encompassing over 300.000 cells across various brain regions and developmental stages. Employing advanced computational methods for data harmonization, cell-type identification, differential gene expression analysis, and inference of cell-cell communication networks, we aimed to identify convergent molecular alterations. Our findings reveal robust, conserved transcriptomic signatures across these diverse models, most notably highlighting widespread dysregulation of excitatory-inhibitory neuronal communication networks and glial dysfunction in astrocytes and mature oligodentrocytes. Crucially, the clinical relevance of these conserved signatures was underscored by significant overlap with known ASD risk genes from the SFARI database and concordance with transcriptomic alterations observed in human postmortem brain studies. Collectively, this integrative study provides a unified framework that surpasses model-specific variability, acknowledging shared molecular mechanisms and cellular pathologies central to ASD.

## METHODS

### Data Acquisition and processing

We curated scRNA-seq data from 11 independent studies investigating animal models of ASD (Table 1). These studies spanned multiple brain regions, developmental timepoints, and genetic or pharmacological ASD models. All data was obtained from public repositories, and if only raw (fasta/fastq) was available, sequencing data was processed through the 10x Genomics CellRanger platform to perform alignment and form corresponding matrix to perform bioinformatic analysis. For each dataset, quality control was performed to remove low-quality cells. In our QC pipeline, we computed key metrics including the log-transformed total RNA counts and the log-transformed number of detected genes and calculated the percentage of mitochondrial gene expression. We applied a custom MAD (Median Absolute Deviation) outlier function (mad_outlier) to flag cells with extreme values (beyond 5 MADs from the median) for the above metrics (Heumos et al., 2023). Cells flagged as outliers were removed from further analysis. Additionally, cells were filtered based on mitochondrial gene content: only cells with less than 10% mitochondrial genes were retained in single-cell datasets, whereas a threshold of 1% was applied for single-nucleus data (Osorio and Cai, 2021).

**Table 1.** List of studies used to construct the integrated object of animal models for ASD.

| DOI | Brain Region | Age | Model | Year | Platform | Methods |
| --- | --- | --- | --- | --- | --- | --- |
| 10.1002/advs.202002117 | Cerebral cortex | P0 | *Sting* Conditional Knockout | 2020 | 10x Genomics | Single-cell RNA sequencing |
| 10.1126/sciadv.abm1077 | Prefrontal cortex and striatum | 3 months | *Setd1a* haploinsufficient | 2022 | 10x Genomics | Single-cell RNA sequencing |
| 10.1371/journal.pgen.1010221 | Cerebral cortex | P5 | *Fmr1* Knockout | 2022 | InDrop | Single-cell RNA sequencing |
| 10.1126/sciadv.ade2441 | Hippocampus | Embryo (17.5 days) | *Foxg1* Knockout | 2023 | 10x Genomics | Single-cell RNA sequencing |
| 10.1016/j.celrep.2023.112712 | Prefrontal cortex | 3-4 months | *Gls1* Conditional Knockout in *CamKIIα*+ neurons | 2023 | GEXSCOPE | Single-nucleus RNA sequencing |
| 10.1016/j.jgg.2023.08.012 | Hippocampus | 3 months | Prenatal exposure to Valproic Acid | 2023 | 10x Genomics | Single-cell RNA sequencing |
| 10.1016/j.celrep.2023.112706 | Cerebral cortex | Embryo (14.5 days) and P0 | *Ube3a* gain-of-function | 2023 | Drop-seq | Single-cell RNA sequencing |
| 10.1093/hmg/ddad155 | Hippocampus | 1 month | *Crlf3* Variant | 2023 | 10x Genomics | Single-cell RNA sequencing |
| 10.1186/s40478-023-01533-w | Cerebral cortex | 3 months | *Slc6a8* Knockout | 2023 | 10x Genomics | Single-cell RNA sequencing |
| 10.1038/s41586-023-05711-7 | Hippocampus | 1 month | NPAS4:NuA4 Conditional Knockout | 2023 | 10x Genomics | Single-nucleus RNA sequencing |
| 10.1371/journal.pgen.1010952 | Cortex and hippocampus | P0 | *Hnrnpu* haploinsufficient | 2023 | 10x Genomics | Single-nucleus RNA sequencing |

### Data Integration, Dimensionality Reduction and Clustering

The raw data were merged and them split by the “paper” identifier to retain sample-specific information. Each individual dataset was normalized using NormalizeData(), and variable features were identified using FindVariableFeatures() with 3000 features. Data were scaled with ScaleData(), and principal component analysis (PCA) was performed on each dataset with 100 PCs computed. These preprocessing steps were applied to all merged datasets. To harmonize data across the 11 studies, we employed a reciprocal principal component analysis (RPCA) integration strategy as implemented in Seurat (v5.0) due to the high number of cells (> 300.000) (Hao et al., 2024). The merged Seurat object was used for integration by invoking the function IntegrateLayers() with the RPCAIntegration method. The original PCA reduction (computed with 100 PCs) was used as input. We next computed a nearest-neighbor graph using the integrated RPCA reduction over the first 100 PCs. Clustering was performed at a high resolution (resolution = 30) to capture a large number of distinct cell populations, with clusters stored in the metadata (e.g., under “rpca_clusters”). For visualization, Uniform Manifold Approximation and Projection (UMAP) was run using the integrated RPCA reduction (using PCs 1–100), which allowed for the detailed exploration of the cellular landscape. This high-resolution clustering strategy facilitated the identification of subtle, cell-type–specific transcriptional signatures that may be critical for understanding the underlying biology.

### Cell-Type Identification

After clustering, cells were further analyzed for differential gene expression. Using FindAllMarkers(), positive markers were identified for each cluster (with a minimum log-fold-change threshold of 0.25). Dimensionality reduction and visualization (via DimPlot()) were performed to assess the distribution of cells by paper and cluster. Hierarchical clustering was conducted in five successive levels and cell-type annotations were assigned by examining the expression profiles of classical gene markers using both literature guidance and data-driven marker identification.

### Differential Expressed Genes Analysis

We also identified differentially expressed genes (DEGs) between ASD and control conditions across all levels of cell classification. DEGs were determined using Seurat’s implementation of the Wilcoxon rank-sum test, applying an average log2 fold change threshold and adjusted p-values to control for multiple testing. Additionally, we compared these DEGs with the enriched genes identified using the FindAllMarkers function for each cluster, to assess whether a given DEG ranks among the most significant genes in that cluster.

### Functional Enrichment and Annotation

Gene Ontology (GO) and KEGG Pathway Analysis: For each cluster, both enrichment genes in each cluster and DEGs between ASD and control samples were subjected to enrichment analysis using standard GO databases (Biological Processes, Molecular Functions, Cellular Components) and KEGG pathways. Gene symbols were then converted to Entrez IDs using the bitr() function from the clusterProfiler package with the mouse annotation database (org.Mm.eg.db). A background gene set was defined using the count data from the RNA assay of a Seurat object. We computed the number of cells in which each gene was expressed (by summing the counts of cells with nonzero expression) and retained only genes with nonzero expression. This background set was similarly annotated with Entrez IDs. For each cluster, GO enrichment analyses were performed separately for the three ontologies: Biological Process (BP), Molecular Function (MF), and Cellular Component (CC). The enrichGO() function was used with the following parameters: the Benjamini–Hochberg (BH) method for p-value adjustment, a p-value cutoff of 0.01, and a q-value cutoff of 0.05. Results were rendered “readable” by converting Entrez IDs back to gene symbols. In parallel, KEGG pathway analysis was conducted using the enrichKEGG() function (with organism set to “mmu” and a p-value cutoff of 0.05) to identify significantly enriched pathways. Enriched terms were visualized using volcano plots and bar charts. IUPHAR and Transcription Factor Classification: DEGs were further categorized based on their classification as transcription factors (TFs) (Hu et al., 2019) or according to physiological and pharmacological criteria using the IUPHAR/BPS Guide to Pharmacology (Armstrong et al., 2020).

### Comparison Between Datasets

To assess the consistency of transcriptomic signatures across the studies included in our integration, we performed several comparative analyses. First, we evaluated the concordance between DEGs identified in the integrated object versus those found within subsets representing each individual reference study. This involved correlation analyses at the second level of cell classification, quantifying agreement using Spearman’s correlation coefficients and statistical significance. Second, we conducted a shared gene analysis among these reference subsets, examining both the number of overlapping DEGs and the consistency of their regulatory direction (up- or downregulation) across studies. Third, to explore cross-model effects, we assessed the expression patterns of key genes associated with the specific genetic perturbations used in the source studies (*Crlf3, Fmr1, Foxg1, Gls1, Hnrnpu, Npas4, Setd1a, Slc6a8, Tmem173*, and *Ube3a*). We specifically examined the regulation of these genes in each subset and within the valproic acid (VPA) pharmacological model subset derived from our integrated object, evaluating if this model significantly recapitulated genetic model-specific alterations at the second level of cell classification. Finally, for a direct comparison with results from an original publication, we selected Donnard et al. (2022), chosen for the compatibility of its reported cell annotations with our integrated dataset. For this specific comparison, neuronal clusters were matched based on our second level of hierarchical classification, while non-neuronal clusters were compared using the more granular third level. DEGs reported in the original paper were filtered (average log2 fold change > 0.25, adjusted p-value < 0.05), and Venn diagrams were generated to visualize the overlap and directional consistency with DEGs identified in our integrated object for corresponding cell types.

### Cell-Cell Communication Analysis

We used CellChat (Jin et al., 2021) to infer and compare cell–cell communication networks across the integrated dataset. The analysis was performed in two stages: Global Analysis: All cell types were considered to construct an overall communication network, where nodes represent cell types and edge thickness indicates the strength of signaling interactions, in the second level on the hierarchical clusterization. Condition-Specific Analysis: Separate networks were constructed for ASD and control conditions both in all clusters in the second level and only in the excitatory and inthibitory neurons in the third level on the hierachical clusterization. Specific signaling pathways (e.g., VEGF, PTN, SLIT, PTPR and SEMA3) were further examined.

### Ligand-Receptor Interaction and NicheNet Analysis

To dissect the molecular underpinnings of altered cell–cell communication between inhibitory (sender) and excitatory (receiver) neurons, we employed NicheNet (Browaeys et al., 2020). This approach predicted the ligand–receptor interactions and downstream target genes that mediate the observed transcriptional changes. Ligand Prioritization: Ligands were ranked based on the area under the precision-recall curve (AUPR) and their fold change between ASD and control conditions. Receptor Analysis: A complementary heatmap was generated to display the expression and predicted interaction strength of key receptors in excitatory neurons. Target Gene Prediction: Predicted target genes in excitatory neurons, as influenced by top inhibitory neuron ligands, were visualized using heatmaps that indicate regulatory potential.

### Validation and Characterization of DEGs with External Databases

To further validate and characterize the DEGs identified between ASD and control conditions (primarily focusing on the third level of hierarchical cell classification), we first assessed their clinical relevance by cross-referencing them with the Simons Foundation Autism Research Initiative (SFARI) Gene database (Abrahams et al., 2013). For each DEG present in the SFARI database, we extracted its associated evidence score and source, comparing these against the gene’s average log2 fold change (avg_log2FC) within each relevant cell cluster. Functional enrichment analysis, including GO and KEGG pathways and classification as a transcription factor or according to physiological/pharmacological criteria using the IUPHAR/BPS was subsequently performed specifically on this subset of SFARI-matched DEGs, following the procedures previously described. Additionally, to investigate patterns of commonality versus cell-type specificity in gene dysregulation, we analyzed the distribution of all DEGs across the third-level clusters. We identified the top 20 most frequently shared DEGs across clusters and the top 10 DEGs uniquely regulated within a single cluster, visualizing their regulation status.

### Comparison with External Human Data

To evaluate the translational relevance of the transcriptomic alterations identified in our integrated animal model dataset, we compared DEG)profiles with those reported in a large human postmortem single-nucleus RNA-seq study of ASD by Gandal et al. (2022). We obtained the published DEG lists from the Gandal et al. study for major cell types across the prefrontal cortex, parietal cortex, and occipital cortex, including gene identity, average log2 fold change, and statistical significance. Cell type annotations were manually matched between our integrated dataset and the Gandal et al. study to ensure comparability across major neuronal and non-neuronal populations. For each matched cell type within each cortical region, we identified the set of common DEGs (shared DEGs) by finding the intersection between the significant DEG lists from our integrated dataset and the corresponding list reported by Gandal et al. (2022). We then calculated the number and percentage of these shared DEGs that exhibited concordant regulation (same direction of avg_log2FC change in both datasets). Furthermore, we computed the Spearman’s correlation coefficient between the avg_log2FC values of the shared DEGs obtained from our integrated dataset and those reported by Gandal et al., determining the statistical significance (p-value) of this correlation. These comparisons were visualized for each cortical region using Venn diagrams to depict DEG counts and overlap, alongside scatter plots illustrating the correlation of avg_log2FC values for shared DEGs, with statistical significance. For shared DEGs in the parietal cortex, we conducted cell-type-specific functional enrichment using SynGO (Koopmans et al., 2019) for synaptic genes in excitatory and inhibitory neurons, and Metascape (Zhou et al., 2019) for mature oligodendrocytes.

### Statistical Analyses

For all differential expression and enrichment analyses, statistical significance was determined using adjusted p-values (e.g., Benjamini–Hochberg correction). Spearman’s correlation coefficients were calculated to evaluate the concordance between DEG sets across datasets and conditions. Graphical representations (e.g., UMAPs, volcano plots, dot plots, heatmaps, and Venn diagrams) were generated in R using ggplot2 and related packages or

## RESULTS

### Characterization of the Integrated Dataset of Animal Models for ASD

We successfully integrated data from 11 studies (Table 1) using advanced integration methods implemented in the Seurat package (*v5.0*). The resulting integrated dataset comprises 155.132 cells from ASD models and 158.075 cells from control animals (Figure 1a). Following data integration, hierarchical clustering was conducted to classify cell types across different functional and transcriptomic levels. At the first level, broad cell groups such as neurons and non-neuronal cells were identified, with the individual signature of the cells becoming more distinct according to functional and transcriptomic characteristics observed in subsequent levels (Supplementary Table 1 and Supplementary Figure 1).

**Figure 1:**
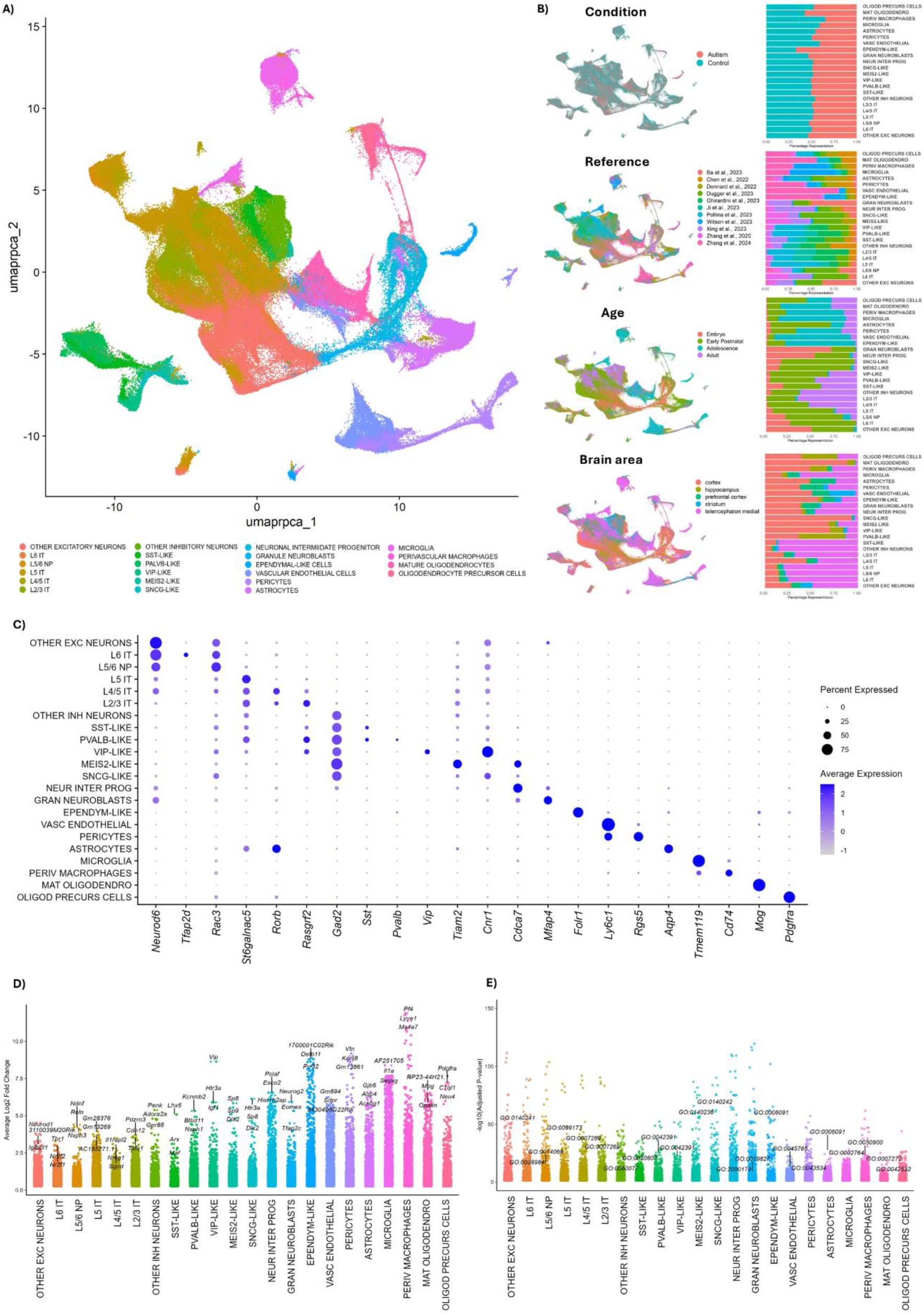
Integration and Analysis of Single-Cell RNA Sequencing from Mice Models of ASD. **A)** UMAP plot displaying clusters after reciprocal PCA (RPCA) integration of single-cell RNA sequencing data from 11 different studies. The plot shows the third level of hierarchical clustering, highlighting distinct cell populations identified across the datasets, colored by major cell types. **B)** Left: UMAPs demonstrate the distribution of cells grouped by metadata categories: condition (ASD vs. control), reference (source of the 11 references used in the integration), age (age of the animals used in the experiments), and brain area (regions of the brain where samples were collected). Right: Corresponding bar plots depict the percentage representation of each feature within each cell identification cluster from Panel A. **C)** Dot plot illustrates the expression levels of key gene markers used to classify each cluster identified at the third level of hierarchical clustering. The dot size represents the proportion of cells expressing the gene, and the color intensity indicates the expression level. **D)** Distribution plot illustrates the top three most differentially expressed genes (average log2 fold change) in each cluster at the third level of hierarchical clustering. **E)** Distribution plot illustrates all significant gene ontology terms and KEGG pathways, for each cluster at the third level of hierarchical clustering. Relevant terms for each cell type function are highlighted in the figure.

Figure 1B shows the distribution of cells across metadata categories. The data demonstrates a uniform distribution of cells across conditions, except for a higher number of cells from ASD-samples for mature oligodendrocytes and ependymal cells. As expected, specific cell types exhibit variation based on experimental design of each reference: progenitor cells are almost exclusive from studies using embryonic or early postnatal animals, for instance, *Ube3a* gain-of-function model represents the highest percentage (38%) of progenitors cells with samples from animals ageing E 14.5 and P0 (Xing et al., 2023), while the lowest percentage of progenitors come from *Slc6a8* knockout model with samples from 3 month old animals (0.33%) (Ghirardini et al., 2023). A complete list of distributions for each reference at every cell classification level is available in Supplemental Table 2. Figure 1C presents a dot plot displaying the expression patterns of canonical marker genes used for cell type annotation at the third hierarchical classification level. Established markers demonstrated high cell-type specificity, such as *Aqp4* for astrocytes and *Vip* for VIP-expressing interneurons. Similarly, *Gad2*, encoding a key enzyme for GABA synthesis, showed the expected high specificity for all identified inhibitory neuron subtypes, including the ‘Other Inhibitory Neurons’ cluster, while being absent in excitatory neurons, confirming its utility as a pan-inhibitory marker. These markers were meticulously selected based on established literature, and a detailed list of markers used at each classification level is provided in Supplementary Table 3.

Figure 1D presents a distribution plot highlighting genes positively enriched in the various cell types by average log2 fold change. The three most enriched genes are emphasized in the figure, often corresponding to genes already known to be associated with the identified cell types, such as *Pdgfra* for oligodendrocyte progenitors and *Mog* for mature oligodendrocytes. A detailed list of the enriched genes at the third level is provided in Supplementary Table 4. To further validate the integration and classification of cell types, we performed a Gene Ontology (GO) analysis for the genes enriched in each cluster. The results are depicted in Figure 1E, where the volcano plot highlights pathways relevant to the functions of each cell type. For instance, in microglia, we highlighted the immune response-regulating signaling pathway (GO:0002764); for vascular endothelial cells, we showed the regulation of angiogenesis pathway (GO:0045765); and for other inhibitory neurons, we highlighted the inhibitory synapse assembly (GO:0060077). The complete list of GO term analyses is available in Supplementary Table 5.

### Transcriptomic alterations between ASD and control

The distribution plot in Figure 2A shows the differentially expressed genes (DEGs) for each cell type, including both upregulated and downregulated genes based on the log2 fold change, comparing ASD to control. The most significantly upregulated or downregulated gene for each type is highlighted in the figure. Notably, genes such as *Ttr* emerge as the most upregulated in several clusters, while *BC023719*, a non-coding RNA is downregulated in 8 clusters, highlighting a potential relevance of those genes across multiple cell types. The complete list of DEGs at the second and third levels is available in Supplementary Table 6.

**Figure 2:**
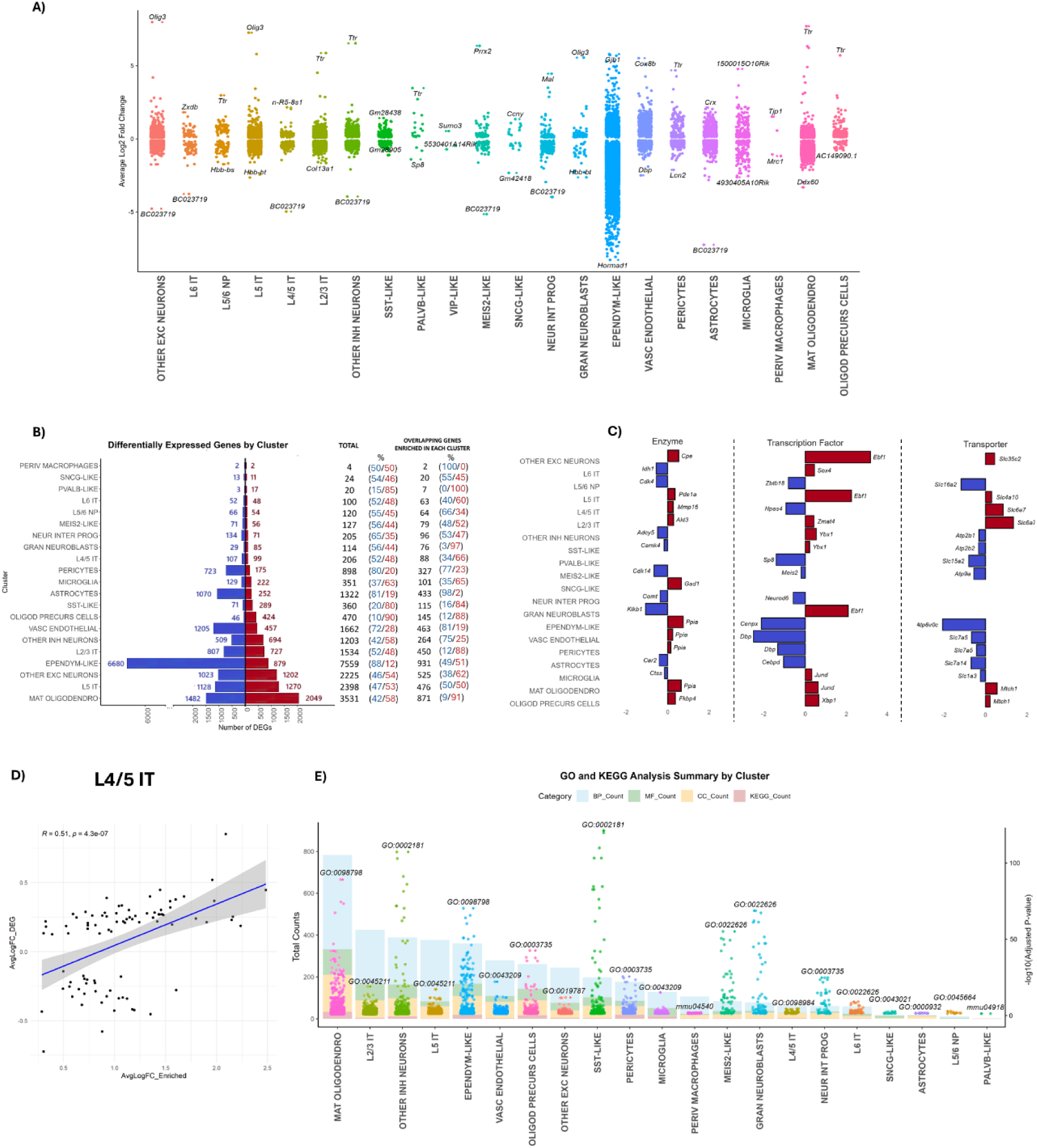
Comparative Analysis Between ASD and Controls in the Integrated Object Third Level. **A)** The distribution plot illustrates the most upregulated and downregulated genes (by average log2 fold change) in the comparison between samples from ASD and controls for each cell type. **B)** The pyramid bar plot shows the burden by clusters with the number of DEGs, ordered by the count of upregulated genes (red) and downregulated genes (blue).The total number of genes is also represented with the percentage of up and downregulated genes, along with the overlap of the enriched genes in each cluster as shown in Figure 1C, representing the percentage of up and downregulated genes again. C) DEGs are represented according to their identity as transcription factors (Hu et al., 2019) or enzymes and transporters based on their physiological or pharmacological classifications using the International Union of Basic and Clinical Pharmacology and British Pharmacological Society (IUPHAR-BPS) (Armstrong et al., 2020). D) An example of Spearman’s correlation analysis between DEGs genes enriched in the cluster L4/5 IT neurons. E) The GO and KEGG analyses by cluster are represented, showing the total counts for each process in the bar plot and each term in the volcano plot, highlighting the most significant term (- log10 adjusted p-value).

The pyramid chart presented in Figure 2B quantifies the DEGs within each cell population, displaying the total counts of upregulated (red bars) and downregulated (blue bars) genes, along with the overall number of DEGs identified per cell type. This visualization effectively highlights the varying transcriptional “burden” associated with the ASD condition across different cellular groups. Ependymal cells, for example, showed the largest absolute number of DEGs (7559), with a strong trend towards downregulation (6680), whereas the perivascular macrophage population exhibited minimal changes (4 DEGs). Additionally, the chart indicates the number of DEGs in each population that also correspond to genes previously identified as significantly enriched markers for that cell type (shown in Figure 1D). This overlap suggests that many dysregulated genes might be integral to the specific functions or identity of these cells. Examining this subset of overlapping DEGs/enriched genes revealed distinct regulatory patterns: in Other Inhibitory Neurons and Astrocytes, 75% and 98% of these overlapping genes were downregulated, respectively. Conversely, in Oligodendrocyte Precursors, 88% of the overlapping genes were upregulated, suggesting potentially different functional consequences of these specific gene expression changes across these cell types.

Further supporting the potential functional relevance of the overlap between DEGs and enriched genes, significant correlations between these gene sets were observed for several cell types (Supplementary Table 7). For example, Figure 2C illustrates a strong positive correlation for L4/5 IT neurons (r = 0.51, p < 0.001); within this specific comparison, *Slc6a7* exhibited the highest positive correlation, while *Camk2n1* showed the most negative correlation. Significant negative correlations were also observed, consistent with findings where key enriched genes were downregulated in the ASD condition. Astrocytes exemplified this pattern, displaying a significant negative correlation (r = −0.17, p < 0.001). Notably, *Aqp4*, identified as the most enriched gene and also known as a key marker for Astrocytes, and crucial for brain water homeostasis, was significantly downregulated in ASD samples within our dataset.

To better understand and characterize the functional roles of the identified DEGs, we categorized them based on annotations TFs or according to their physiological and pharmacological classifications within the IUPHAR/BPS Guide. Figure 2D illustrates representative significant DEGs across several key functional categories (such as enzymes, transporters and TFs) for each cluster, with the complete list of categorized DEGs available in Supplementary Table 8. Notable examples include alterations within the enzyme category, such as the downregulation of *Comt* in Neuronal Intermediate Progenitors and the upregulation of *Gad1* in SNCG-like interneurons, suggesting potential impacts on catecholaminergic and GABAergic neurotransmission, respectively. Among transcription factors, *Ebf1* showed significant upregulation in Other Excitatory Neurons, L5 Neurons and Granule Neuroblasts. Furthermore, widespread alterations were identified across various members of the Solute Carrier (SLC) family of transporters within both neuronal and non-neuronal populations, indicating potential disruptions in fundamental physiological transport processes within the ASD models.

We subsequently performed GO and KEGG pathway enrichment analyses on the DEGs from each cluster, summarized in Figure 2E. This visualization depicts the total count of enriched terms across GO categories and KEGG pathways, with point attributes reflecting statistical significance. While no KEGG pathways directly mapping to ASD as a disease were identified, several GO terms pertinent to brain function were significantly enriched across multiple cell types. Key processes included ‘cognition’ (GO:0050890), found enriched in diverse populations such as various excitatory/inhibitory neurons (e.g., L2/3 IT, L5 IT, SST-like) and glial cells (e.g., microglia, astrocytes, oligodendrocytes), and ‘learning’ (GO:0007612), noted particularly in excitatory neuron subtypes (e.g., Other Excitatory, L4/5 IT). These results highlight potential alterations in critical neural pathways. Interestingly, a high absolute number of DEGs in certain cell types, such as ependymal cells, did not necessarily correspond to a high number of significantly enriched functional terms based on GO analysis. This observation suggests that in some cellular contexts, widespread gene expression changes might be distributed across many pathways below significance thresholds, or that the functions of many DEGs remain poorly characterized. The complete list of enriched GO and KEGG terms for all clusters is detailed in Supplementary Table 9.

### Comparison Between Integrated Object and Individual Datasets

To investigate the transcriptomic signatures across studies used in our integration and integrated object, we conducted analyses to compare conserved dysregulation patterns while highlighting dataset-specific nuances. In Figure 3A, correlation analysis revealed varying degrees of concordance between DEGs in the integrated dataset and the subset of individual references in our cell type classification at the second level. Strong correlations were observed for inhibitory/excitatory neurons and progenitor cells across multiple references, suggesting these cell types may exhibit conserved transcriptomic dysregulation across datasets. Despite nearly every reference showing a significant correlation (Supplementary Table 10), lower correlations values in some clusters or references underscore potential dataset-specific variations.

**Figure 3:**
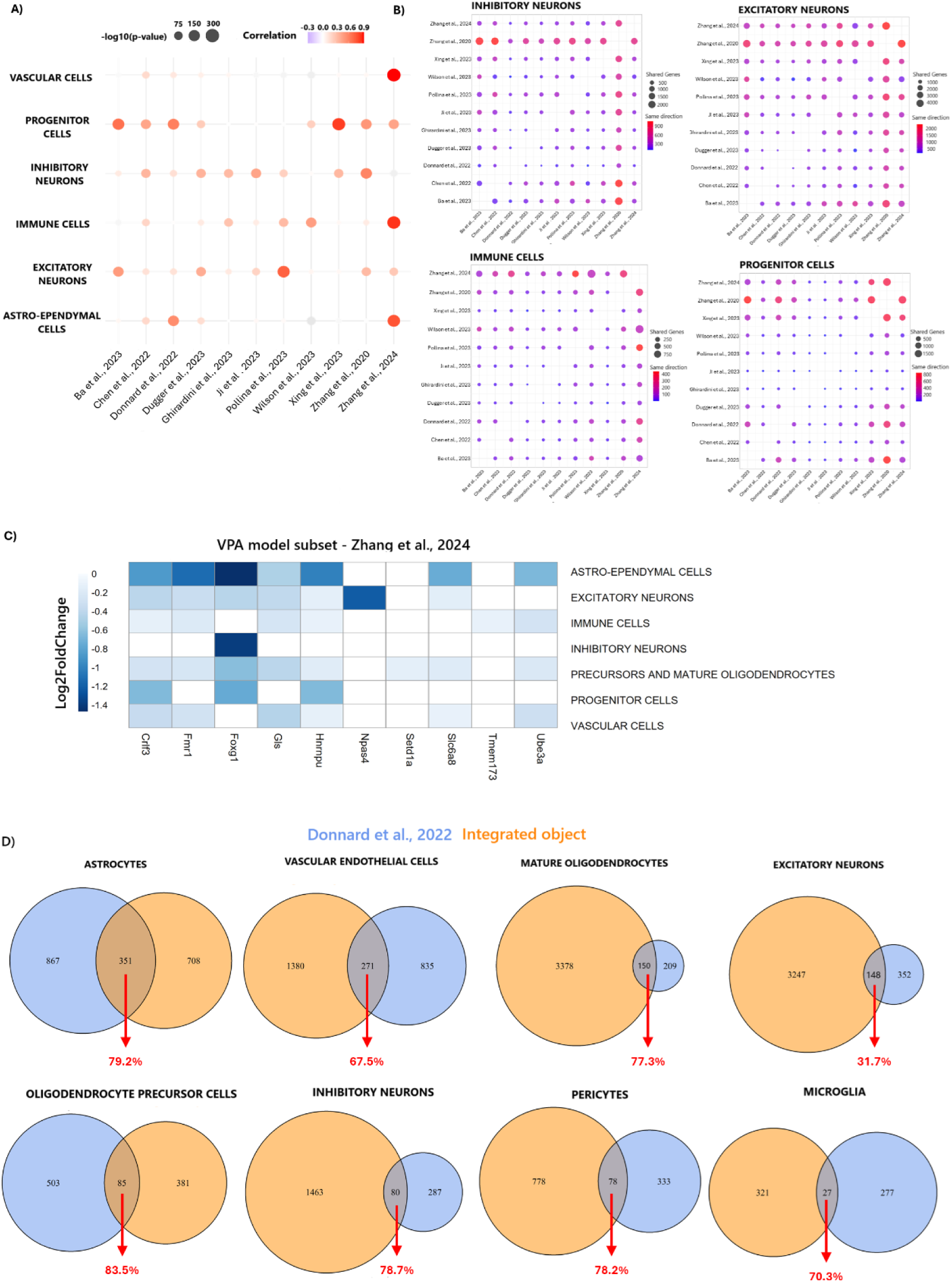
Comparative Analysis of Differentially Expressed Genes Across Integrated Object and Individual Datasets in the Single-Cell ASD Study. **A)** Correlation analysis of DEGs between the integrated dataset and subsets from individual reference studies included in the integration. Each dot represents a cellular cluster, with dot size indicating statistical significance (-log10(p-value)) and color intensity reflecting the strength of the correlation (scale: −0.3 to 0.9). **B)** Shared gene analysis among each reference study subset in the integrated object, focusing on specific cell types: inhibitory neurons, excitatory neurons, immune cells, and progenitor cells. Dot size represents the number of shared DEGs, while color indicates the proportion of shared genes exhibiting consistent regulation (up- or downregulated) across datasets. **C)** Heatmap showing the log2FC of modulated genes in each genetic paradigm model for ASD in the subset of the pharmacological model of valproic acid (VPA) subset in our integrated object. **D)** Venn diagrams compare DEGs from the integrated dataset with those found in the selected original reference study (Donnard et al., 2022). DEGs from the Donnard dataset were filtered for both statistical significance (adj_pvalue < 0.05) and an average log2 fold change > 0.25, ensuring robust comparisons. Each diagram illustrates the overlap of DEGs for specific cellular clusters and red numbers indicating the % of shared DEGs that are regulated in the same direction

Furthermore, the shared DEG analysis among subsets of each reference study, as shown in Figure 3B and drawn from the integrated dataset, highlights the close alignment of these subsets in certain clusters. The degree of overlap and the consistency in regulatory direction (up- or down-regulated) varied based on the reference and cell type analyzed. Some subsets demonstrated stronger overlap and higher consistency, while others exhibited more divergence. For instance, inhibitory and excitatory neurons tended to share a greater number of genes (and DEGs in general) with consistent regulation across references, whereas immune cells and progenitors showed less overlap, likely related to the differing ages of subjects in each reference. Despite these differences, all subsets shared at least some DEGs with consistent regulatory direction, underscoring the presence of conserved patterns across studies for specific cell types.

To assess consistency across different ASD models, we examined the expression of genes specifically targeted or modulated in the source studies within their corresponding cell subsets in our integrated dataset. For instance, *Gls1*, knocked out in *CamKIIα*+ neurons by Ji et al. (2023), displayed concordant downregulation in excitatory neurons in our integrated analysis and in datasets from Chen et al. (2022), Zhang et al. (2024), and Pollina et al. (2023) (Supplementary Figure 3). Similarly, *Npas4*, knocked out by Pollina et al. (2023), was significantly downregulated in the corresponding excitatory neuron subset of our integrated data and also showed reduced expression in subset data from Ji et al. (2023), Ghirardini et al. (2023), and Zhang et al. (2024). This cross-model consistency supports the polygenic nature of ASD, highlighting how multiple gene expression changes contribute to the phenotype. The observation that certain genes are consistently altered across various models points to their potential importance in ASD etiology within specific cell types.

We extended our analysis to the VPA pharmacological model subset (Zhang et al., 2024), focusing on those key ASD-associated genes in the genetic models. Strikingly, whenever these genes showed significant differential expression compared to controls within this VPA subset, the change was always downregulation (Figure 3C). For instance, *Fmr1* and *Foxg1* were downregulated across most identified cell types, although *Foxg1* downregulation was the only gene regulated in inhibitory neurons among these examples. Additionally, *Tmem173* (STING) downregulation was restricted to immune cells, consistent with the subset of source study (Zhang et al., 2020). These findings underscore the utility of the VPA model for ASD research, demonstrating its capacity to alter the expression of multiple relevant genes across diverse cell populations.

We also proposed a new analysis to compare the DEGs identified in a specific original reference dataset with our integrated dataset without creating subsets. Venn diagrams illustrate the overlap of DEGs for specific clusters between the original Donnard et al., 2022 dataset and our integrated dataset in Figure 3C. For instance, in astrocytes, 351 shared DEGs were identified, with 278 showing consistent regulation (79.2%). Similarly, vascular endothelial cells and mature oligodendrocytes exhibited 67.5% and 77.3% consistent DEGs, respectively. Clusters with smaller overlaps, such as microglia (27 DEGs, 70.3% consistent) and pericytes (78 DEGs, 78.2% consistent), highlight the complexity of cell-type-specific transcriptional regulation and potential variations in experimental approaches or sample characteristics between the integrated dataset and the Donnard et al., 2022. Despite these variations, the presence of consistently regulated DEGs across all analyzed clusters indicates that key biological signals are preserved between the datasets.

### Investigating Cell-Cell Communication Disturbances

Analysis of overall cell communication across all cell types in the integrated dataset, regardless of condition, revealed a highly interconnected network (Supplemental Figure 2A). Key interactions were observed among both neuronal and non-neuronal cell types, reflecting the robust integration and accurate annotation of the dataset. The VEGF signaling pathway was analyzed to validate the cell type annotations (Supplemental Figure 2B). This pathway specifically involved interactions between vascular and astro-ependymal cells, which align with their known physiological roles in maintaining vascular and neural homeostasis, further supporting the accuracy of the annotations. When comparing ASD to control conditions, cellular communication at the second annotation level revealed a significant increase in interactions between inhibitory and excitatory neurons in ASD (Figure 4A).

**Figure 4:**
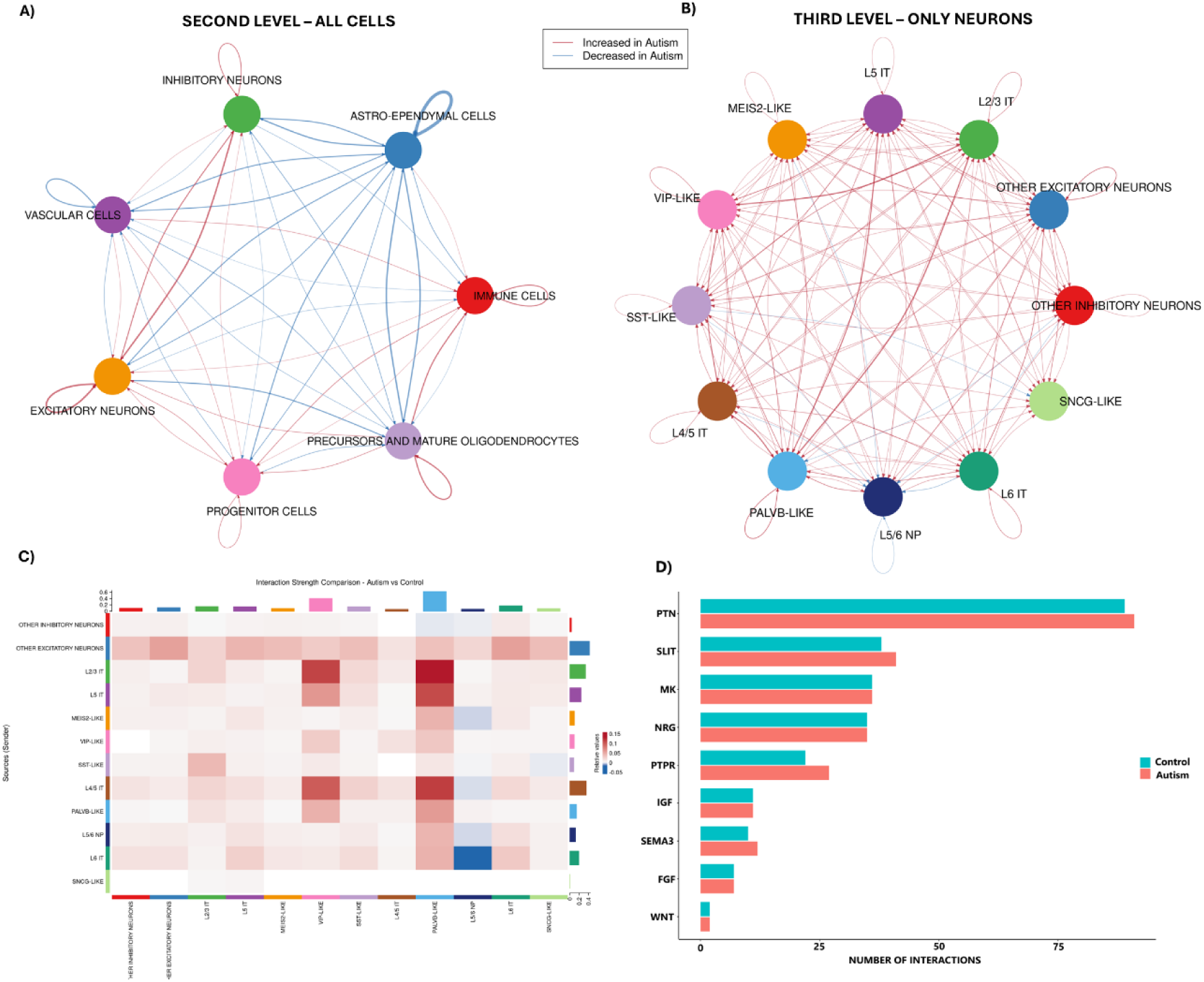
CellChat Analysis of Cellular Communication in Condition-Specific Comparisons (ASD vs. Control). **A)** Comparison of cellular communication networks between ASD and control samples at the second annotation level, including all cell types, shows increased communication in ASD (red edges) between inhibitory and excitatory neurons. **B)** Interactions at the third annotation level, focusing only on neuronal subtypes, reveal that this increase in communication is consistent across most neuronal subtypes. **C)** Interaction strength comparison heatmap illustrating the differences in signaling strength between ASD and control across all neuronal subtypes at the third annotation level. Red indicates stronger signaling in ASD, while blue represents stronger signaling in control. **D)** Bar plot showing the number of interactions for key signaling pathways between ASD and control. Pathways such as PTN, SLIT, and NRG exhibit increased interactions in ASD.

This increase suggests a general dysregulation in neuronal signaling that could underlie the hyperconnectivity often reported in ASD-related phenotypes. To further investigate this heightened neuronal communication, a more detailed analysis was conducted at the third annotation level, focusing exclusively on neuronal subtypes (Figure 4B). The results demonstrate that this increase in communication is not limited to specific neuronal subtypes but is instead a broad effect across most neuronal populations. Subtypes such as L4/5 IT and PALVB-like neurons showed substantial increases in interaction strength in ASD compared to control, as illustrated in the heatmap (Figure 4C), highlighting the widespread dysregulation of neuronal networks.

To identify the signaling pathways driving these differences, we analyzed the key signaling pathways that exhibited altered activity in ASD (Figure 4D). Pathways such as PTN, SLIT, PTPR, and SEMA3 showed a significant increase in the number of interactions in ASD, indicating their potential role in mediating the increased neuronal communication. In contrast, pathways like WNT and FGF displayed more balanced activity, suggesting that some pathways may remain relatively unaffected. Further analysis of signaling dynamics revealed distinct patterns of incoming and outgoing signals for neuronal subtypes in ASD compared to control. Incoming signaling patterns (receiver signals) highlighted a global increase in pathway activity across neuronal subtypes in ASD, with particularly strong signals observed for the PTN and SLIT pathways (Supplemental Figure 2C). Visual inspection confirms this widespread increase, showing heightened receiver strength for PTN and SLIT across nearly all analyzed excitatory (e.g., L2/3 IT, L5 IT, L4/5 IT) and inhibitory (e.g., VIP-LIKE, SST-LIKE, PVALB-LIKE) subtypes in ASD compared to controls. PTPR and SEMA3 also show increased reception, though perhaps slightly more concentrated in IT and L5/6 NP subtypes. Outgoing signaling patterns (sender signals) mirrored this trend, with ASD showing increased signaling activity originating from key neuronal subtypes (Supplemental Figure 2D). Specifically, for the PTN, PTPR, and SEMA3 pathways, this enhanced sender activity appears largely driven by excitatory clusters such as L2/3 IT, L5 IT, L4/5 IT, and L5/6 NP. The SLIT pathway shows a similar pattern of increased sending from these excitatory neurons, but also potentially increased contributions from various inhibitory subtypes in ASD. These findings collectively suggest that ASD is characterized by widespread alterations in neuronal communication, driven by increased activity in key signaling pathways and dysregulated sender-receiver dynamics.

Building on the CellChat analysis, which showed increased communication between inhibitory and excitatory neurons in ASD, we aimed to investigate the molecular mechanisms driving these changes in more detail. Given the key role of inhibitory and excitatory neurons in maintaining the balance of neuronal network activity, we concentrated on inhibitory neurons as senders and excitatory neurons as receivers, as shown in Figure 5. This choice was influenced by the observation that inhibitory neurons frequently play a role in shaping and modulating downstream signaling in neural circuits, and their dysregulation could lead to cascading effects on excitatory neurons, potentially contributing to the extensive communication changes seen in ASD.

**Figure 5:**
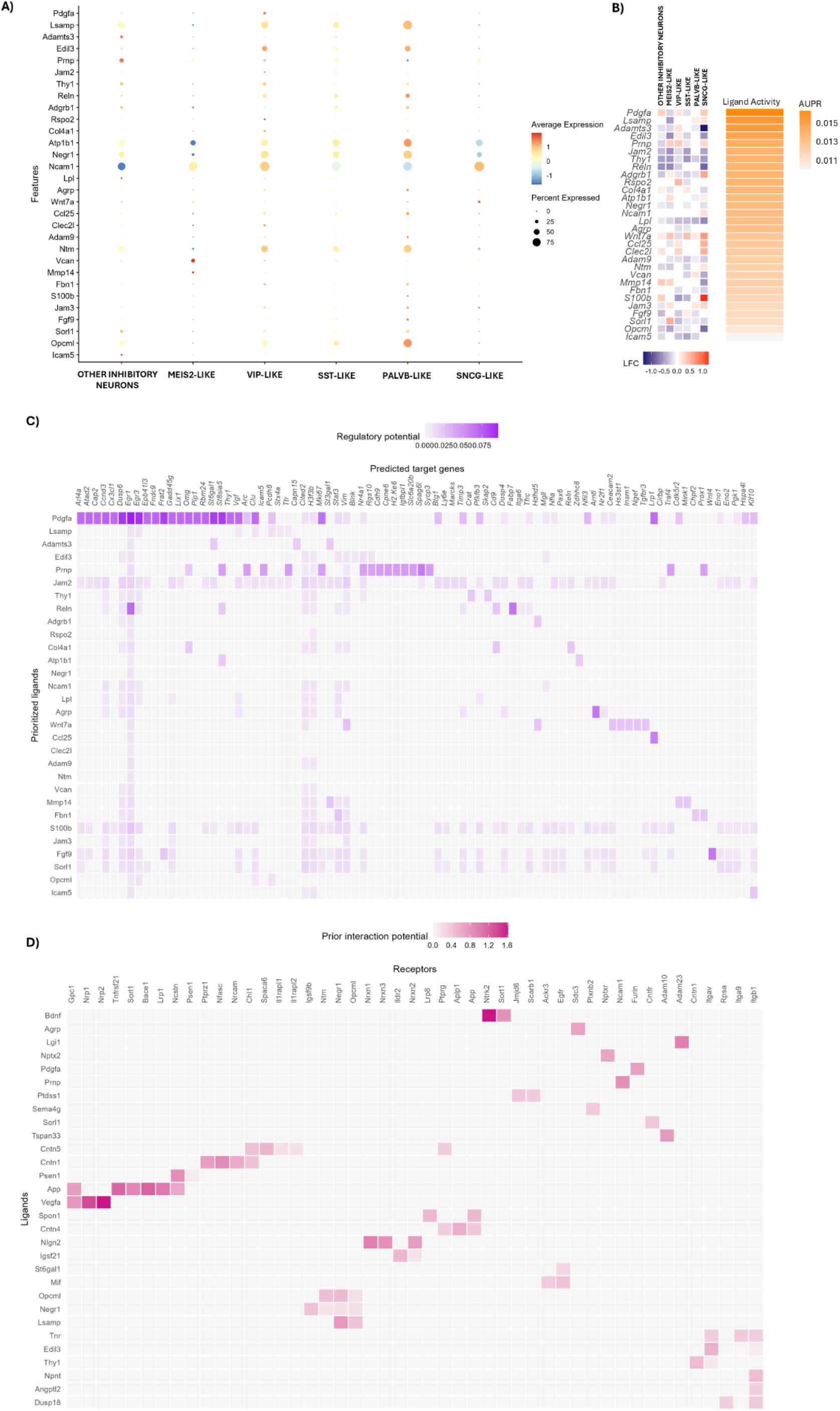
NicheNet Analysis of Ligand-Receptor Interactions Between Inhibitory Neurons (Senders) and Excitatory Neurons (Receivers). **A)** Dot plot displaying the expression levels (average expression) and percentage of cells expressing the top ligands identified in inhibitory neurons, categorized by neuronal subtypes. Each dot’s size indicates the proportion of expressing cells, while the color denotes the average expression level. **B)** Heatmap ranking the top ligands based on the area under the precision-recall curve (AUPR), their log fold change (LFC) in ASD compared to control, and their expression across inhibitory neuron subtypes. **C)** Heatmap illustrates the predicted target genes in excitatory neurons (receivers) affected by the top ligands. The intensity of purple shading represents the regulatory potential of each ligand-target gene interaction, reflecting the anticipated impact of ligand activity on target gene expression in the receiver cells. **D)** Heatmap of the top receptors in excitatory neurons, displaying the prior interaction potential for each ligand-receptor pair. The intensity of shading reflects the strength of the predicted interaction, emphasizing key ligand-receptor interactions facilitating communication between inhibitory and excitatory neurons.

The expression levels and distribution of the top ligands identified in inhibitory neurons are illustrated in Figure 5A. Ligands such as *Pdgfa*, *Lsamp*, and *Adamts3* exhibited high ligand activity and were broadly expressed across multiple inhibitory neuron subtypes. Other ligands, including *Wnt7a* and *Jam2*, displayed more restricted expressions, suggesting potential roles in subtype-specific signaling. These findings highlight the diverse signaling repertoire of inhibitory neurons, which may contribute to the observed communication dysregulation in ASD.

To quantify ligand activity and assess their differential regulation in ASD, we ranked the top ligands based on their area under the precision-recall curve (AUPR) and analyzed their fold change between ASD and control conditions (Figure 5B). Although the AUPR values are relatively low, indicating limited statistical power to predict condition-specific DEGs (which is expected given the complex etiology of ASD), ligands such as *Pdgfa*, *Adamts3*, and *Lsamp* emerged as the most active. These ligands showed significant regulation across multiple subtypes of inhibitory neurons, highlighting their potential roles in ASD-related signaling. In contrast, ligands such as *Jam2*, *Thy1*, and *Lpl* exhibited consistent downregulation across all inhibitory neuron subtypes, suggesting a more global pattern of dysregulation that may broadly impact neuronal communication in ASD.

Next, we examined the predicted target genes in excitatory neurons affected by the top ligands (Figure 5C). Notably, we found in the the primary targets of *Pdgfa*, *Arl4a* and *Atad2*, exhibited opposite regulation, as observed in the complete DEGs list (Supplementary Table 6): *Arl4a* was consistently upregulated in L5 IT neurons and other excitatory subpopulations, while *Atad2* was downregulated in these groups. In contrast, *Egr1*, a target gene with high regulatory potential for the ligand *Reln*, was significantly downregulated in both L4/5 IT and L5 IT neurons. These findings suggest that dysregulated inhibitory neuron activity in ASD may lead to widespread alterations in the gene expression profiles of excitatory neurons, potentially amplifying communication changes and contributing to network-level dysregulation.

Next, we investigated ligand-receptor interactions to identify the key receptors mediating these signaling changes (Figure 5D). In this analysis, ligands with a higher potential for interaction with specific receptors were prioritized. When we checked the top ligands expression when comparing ASD and control (Supplementary Table 6), *Bdnf* was particularly notable: while its receptor *Ntrk2* showed no significant changes, the receptor *Sort1* was significantly downregulated in L5 IT neurons. Additionally, other key receptors such as *Nrp1* and *Nrp2* were identified. *Nrp2* was consistently upregulated in L5 IT neurons, whereas *Nrp1* displayed a mixed regulatory pattern, being downregulated in some excitatory subtypes (e.g., Other Excitatory and L5/6 NP neurons) and upregulated in others (e.g., L2/3 IT and L5 IT neurons). This mixed pattern suggests that other ligands or mechanisms may influence receptor activity, pointing to subtype-specific responses or distinct functional roles in excitatory neurons.

### Comparison between integrated datasets and clinical evidence

Lastly, we propose comparing our findings with existing literature detailing alterations associated with ASD in humans and mouse models os ASD. First, we conducted an analysis comparing the differentially expressed genes (DEGs) identified in the third level of cell annotations with the SFARI database (Abrahams et al., 2013). This database serves as a comprehensive resource that curates genes linked to ASD based on evidence from genetic, functional, and clinical studies. Each gene is assigned a score from 1 to 3, where a score of 1 indicates the strongest evidence for ASD association, a score of 2 denotes strong but secondary evidence, and a score of 3 implies emerging or preliminary evidence. This analysis enabled us to assess how well the genes identified in our integrated dataset associate with known ASD linked genes, validating our findings and highlighting potential novel targets (Figure 6).

**Figure 6:**
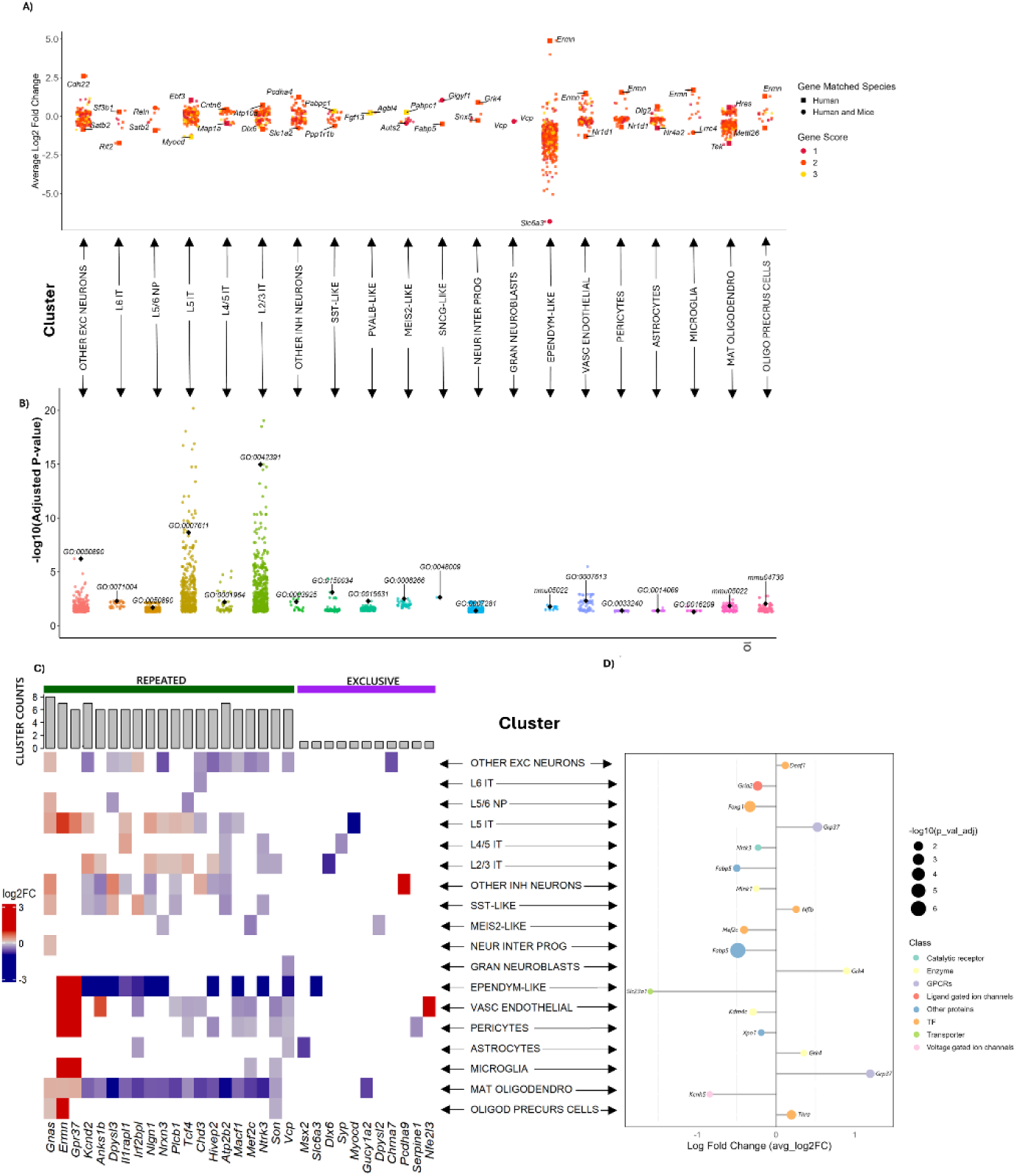
Matching DEGs with the SFARI Database and Functional Analysis. **A)** The scatter plot illustrates the average log2 fold change (avg_log2FC) of DEGs per cluster matched to the SFARI database. Points are color-coded based on gene score (1-3), and shape-coded by species evidence (square for humans or circle for both human and mouse). Genes with higher scores (1 or 2) indicate greater confidence as ASD-associated genes. **B)** Gene ontology analysis of matched DEGs displays the top enriched biological processes for each cluster. Points represent the significance (-log10 adjusted p-value) of the GO terms, with certain significant terms related to ASD being highlighted. **C)** The top 20 most shared and the top 10 unique DEGs across clusters by absolute log2FC, with bars representing the number of clusters in which each gene is either upregulated or downregulated in the heatmap, by avg_log2FC (−3, 3). **d)** A lolliplot categorizes the top DEG (by avg_log2FC) for each cluster based on IUPHAR or TF classification. Categories include catalytic receptors, enzymes, transporters, transcription factors, and others, highlighting their functional roles.

Most matched genes in the SFARI database were high-confidence ASD-associated genes, with scores of 1 or 2, indicating strong evidence (Figure 6A). Notably, when analyzing the proportion of DEGs within each cell population that matched SFARI, we observed that specific neuronal subtypes exhibited some of the highest percentages of matches. Specifically, clusters such as L4/5 IT (16.5%), L2/3 IT (11.5%), L5/6 NP (10.8%), and Other Inhibitory Neurons (10.6%) were among those with the highest proportion of SFARI genes among their respective DEGs, along with non-neuronal populations like Pericytes (10.1%). Illustrating this overlap, known ASD-associated genes such as *Ermn*, supported by evidence from human and/or mouse studies, was identified as a DEG with highest average log2 fold changes and upregulted in multiple non-neurons cell types (Microglia, Vascular Endothelial Cells, Mature Oligodendrocytes, Pericytes, Oligodendrocyte Precursor Cells, Ependymal-Like Cells). The complete list of SFARI-matched DEGs per cluster is detailed in Supplementary Table 11. Furthermore, although many matches were identified, many DEGs from our analysis remain unmatched or lack evidence for ASD association, representing potential novel candidates for future investigation.

Moreover, the GO and KEGG analysis of matched DEGs revealed significant enrichment terms associated with ASD across clusters (Figure 6B and Supplementary Table 12). In neuronal clusters, processes such as cognition (*GO:0050890*; significantly enriched in Palvb-Like, L5-6 NP, L2-3 IT, Other Excitatory Neurons, L5 IT), learning or memory (*GO:0007611*; enriched in Palvb-Like, L5-6 NP, L2-3 IT, Other Excitatory Neurons, L5 IT), and regulation of membrane potential (GO:0042391; enriched in Palvb-Like, L5-6 NP, L2-3 IT, Other Excitatory Neurons, L5 IT) were particularly enriched, aligning with core ASD-related phenotypes. In non-neuronal clusters, processes related to antioxidant activity (*GO:0016209*; enriched in Microglia) and pathways of neurodegeneration (KEGG:mmu05022; enriched in Ependymal-Like Cells, Mature Oligodendrocytes) were significantly enriched, suggesting broader systemic contributions to ASD pathology. These findings not only validate the relevance of the identified DEGs to ASD but also provide valuable insights into the functional pathways altered across diverse cell types, offering potential for further exploration.

We observed that many matched DEGs were shared across clusters, and we illustrate some of these shared genes (Figure 6C). Genes such as *Kcnd2* and *Tcf4* exhibited complex regulatory patterns, being upregulated in some cell types and downregulated in others, indicating intricate, cell-type-specific roles. In contrast to these shared DEGs, we also identified genes uniquely regulated in specific clusters. For instance, *Chrna7* was uniquely downregulated in other excitatory neurons, while *Syp* and *Pcdha9* were uniquely upregulated in other L5-IT and inhibitory neurons, respectively (Figure 6B, right panel). These findings suggest that while some genes have broad, complex effects across cell types, others serve specialized roles within specific cellular contexts.

We categorized the top DEGs by p_val_adjusted on each cluster using IUPHAR and TF annotations (Figure 6D) to further characterize them. Notably, *Gria2*, a gene coding a glutamate receptor subunit essential for synaptic plasticity, was identified as the top down-regulated gene in L6 IT excitatory neurons, emphasizing its role in ASD-related synaptic dysregulation. Similarly, *Mef2c*, a transcription factor critical for neurodevelopment, emerged as the top downregulated gene in MEIS-2 interneurons. Other genes significantly linked to ASD, such as *Deaf1* (upregulated in Other Excitatory Neurons) and *Foxg1* (downregulated in L5-6 NP), were also highlighted. These transcription factors are known to be associated with syndromic forms of ASD, where mutations in these genes lead to severe developmental disorders, further underscoring their relevance to neuronal communication and ASD-related mechanisms.

Building on the comparison with the SFARI database, we sought to validate the relevance of the DEGs identified in our integrated object by comparing them to single-cell human post-mortem data. To achieve this, we curated literature data, selecting the dataset with the highest number of cells available (over 250,000 nuclei from six individuals with ASD exhibiting strong differential expression signatures and six matched control subjects) from Gandal et al., 2022. This study conducted transcriptomic profiling across the prefrontal, parietal, and occipital cortices of individuals with ASD, identifying significant dysregulation across multiple cortical regions. By comparing our integrated DEGs with those reported in their study, we aimed to pinpoint shared transcriptional changes and evaluate the overlap in cell-type-specific dysregulation (Figure 7 and Supplementary Table 13).

**Figure 7:**
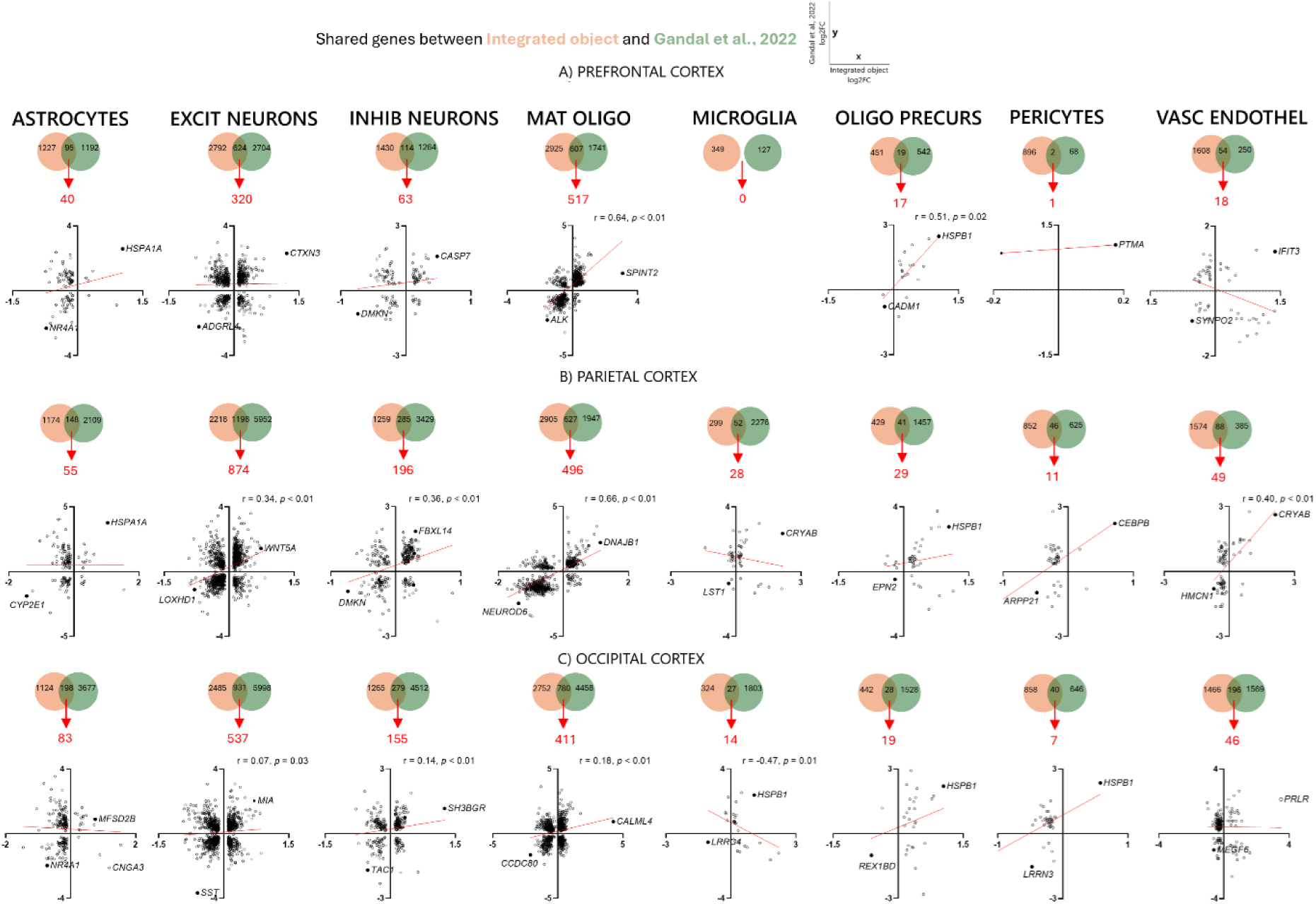
Shared DEGs in common cell types between the integrated object and cortical transcriptomic data from Gandal et al. (2022). Venn diagrams show the total number of DEGs identified in each dataset for specific cell types, with intersections representing shared DEGs and red numbers indicating those that are regulated in the same direction. Asterisks above the Venn diagrams highlight clusters with significant correlations between datasets (*p* < 0.05). Correlation plots below illustrate the avg_log2FC (y axis for Gandal et al., 2022 and x axis for our integrated object), the for shared DEGs on each cell type**: A)** Prefrontal cortex (PFC), **B)** Parietal cortex, and **C)** Occipital cortex.

In the prefrontal cortex, excitatory neurons shared 624 DEGs, of which 320 were regulated in the same direction (Figure 7A) with *CTXN3* gene consistent upregulated while *ADGRL4* consistent downregulated. Interestingly, mature oligodendrocytes exhibited 607 shared DEGs, with 517 consistently regulated with a significant correlation. Oligodendrocyte precursor cells also showed a significant correlation despite having only 19 shared genes. Other clusters in the PFC revealed minimal overlap, with microglia sharing no genes and pericytes only one. In the parietal cortex, excitatory neurons had the highest overlap with 1198 shared DEGs, 874 of which were significant consistently regulated, with a representant of the WNT signialing, the *WNT5A* gene being upregulated in both datasets. Inhibitory neurons and mature oligodendrocytes demonstrated notable overlaps, with 59% and 79% of shared DEGs conserved, respectively, and significant correlations. Vascular endothelial cells also showed substantial overlap with 49 of 88 shared genes consistently regulated (Figure 7B). In contrast, the occipital cortex exhibited the least consistent expression patterns relative to the integrated dataset (Figure 7C). Although excitatory neurons in this region had 931 overlapping DEGs, the correlation was weak, and mature oligodendrocytes showed only a modest correlation. Notably, microglia in the occipital cortex demonstrated a significant negative correlation.

Overall, These findings position the parietal cortex as the region with the highest number of shared genes across clusters and the most consistent regulation compared to our integrated object, while the occipital cortex exhibited the least consistent expression patterns compared to the integrated object, which was expected since this cortical region was not included in the samples used for the integration. Given the results, we further investigated the shared DEGs regulated in the same direction between the integrated object and the parietal cortex DEGs in clusters with significant and consistent correlations, specifically, excitatory neurons, inhibitory neurons, and mature oligodendrocytes. For this analysis, we focused on identifying enriched terms related to functional pathways within these shared genes. To explore synaptic-related processes in excitatory and inhibitory neurons, we used SynGO, a publicly available knowledge base for synapse research that provides a detailed ontology for synaptic components and functions (Figure 8A and B). For mature oligodendrocytes, we utilized Metascape, a comprehensive web-based platform for functional enrichment, interactome analysis, and gene annotation (Zhou et al., 2019), which offers a holistic approach to identifying enriched pathways (Figure 8C and D).

**Figure 8:**
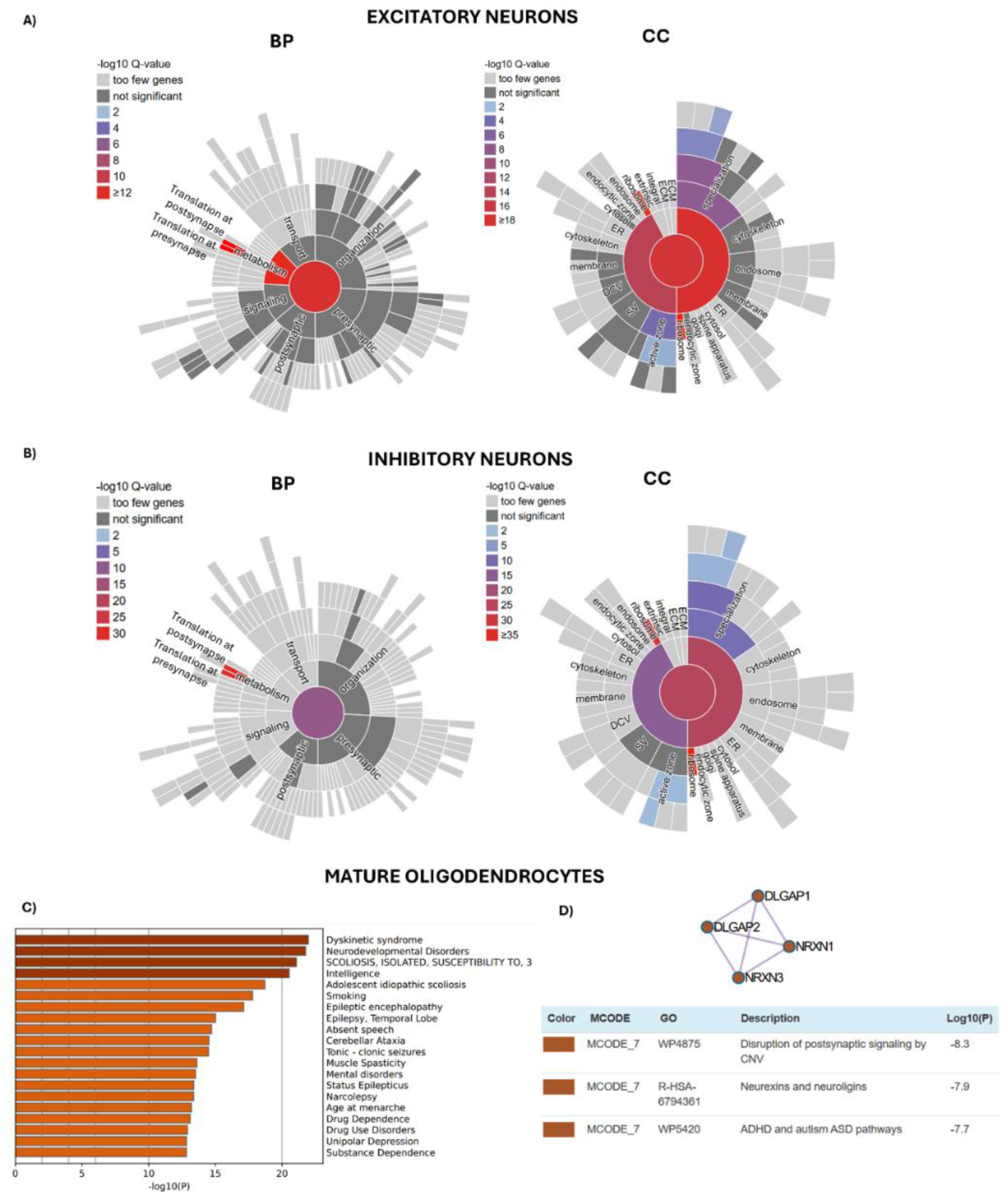
Functional Enrichment Analysis of Shared DEGs Between Integrated Object and Parietal Cortex. SynGO enrichment analysis shows significant biological processes (BP, left) and cellular components (CC, right). The color gradient indicates the significance level (-log10 Q-value), with darker shades representing greater significance for **A)** Excitatory neurons and **B)** Inhibitory neurons. **C)** Metascape functional enrichment analysis for mature oligodendrocytes displays the top enriched human diseases related to the associated genes. The bar plot illustrates significance (-log10 P-value). **D)** Metascape network analysis for mature oligodendrocytes identifies key regulatory genes (*DLGAP1, DLGAP2, NRXN1,* and *NRXN3*) within enriched pathways. The nodes indicate genes, while edges represent shared pathway membership.

For both excitatory and inhibitory neurons, enrichment analysis using SynGO revealed significant biological processes (BPs) and cellular components (CC) associated with synaptic activity (Figure 8A and B). This enrichment in synaptic processes was exemplified by key differentially expressed genes: in Excitatory Neurons, presynaptic functions involved genes such as *CACNA1A* (essential for the Ca2+ influx release) and *UNC13A* (crucial for vesicle priming), while postsynaptic processes were represented by *GRIN2B* (NMDA receptor subunit), CACNG2 (AMPA receptor regulation), and the signaling kinase *CAMK2A*. In inhibitory neurons, presynaptic enrichment highlighted genes such as *ATP6V0A1* (fundamental for neurotransmitter loading into vesicles) and *KCNMA1* (release modulator), while postsynaptic processes involved genes like the signaling kinase *CAMK2A* and the signaling molecule *NRG1*. Furthermore, BP analysis at the second level highlighted important pathways such as “translation at presynapse” and “translation at postsynapse” in both neuronal types. This was particularly evident in inhibitory neurons, where the presynaptic list showed strong enrichment in numerous ribosomal protein genes (e.g., multiple *RPL* and *RPS* genes), emphasizing the critical roles of synaptic signaling and local protein synthesis in ASD-related dysregulation. The complete list of SynGO analysis is available in Supplementary Table 14.

For mature oligodendrocytes, Metascape analysis using the DisGeNET platform (Piñero et al., 2017) revealed significant enrichment of human disease-associated genes among the shared DEGs (Figure 8C). Pathways such as “Dyskinetic Syndrome,” “Neurodevelopmental Disorders,” “Epilepsy,” and “Mental Disorders” were notably enriched, suggesting that the conserved DEGs identified in animal models and human clinical studies implicate oligodendrocytes as direct contributors to ASD pathology. Importantly, when examining the protein-protein interaction (PPI) network, a functional module containing *DLGAP1, DLGAP2, NRXN1*, and *NRXN3* emerged as a key regulatory hub (Figure 8D). This module was enriched in pathways such as “Disruption of Postsynaptic Signaling by CNV” (WP4875), “Neurexins and Neuroligins” (R-HSA-6794361), and “ADHD and ASD Pathways” (WP420), further reinforcing the role of oligodendrocytes in modulating neuronal connectivity. The complete list of Metascape analysis is available in Supplementary Table 15.

## DISCUSSION

By systematically integrating 11 scRNA-seq datasets from multiple ASD models, we constructed a unified single-cell reference for ASD-related transcriptional dysregulation. This approach allowed us to harmonize cell-type annotations across studies and identify shared molecular signatures of ASD at a cellular level that had not been previously achieved. The integration was meticulously curated to avoid overcorrection while preserving biologically significant cell-type distinctions. One of the key advantages of this method was the ability to compare various genetic and environmental ASD models within a standardized framework, revealing consistent alterations across diverse experimental paradigms. While this approach has previously been successfully utilized in the context of renal and metabolic disorders (Hrovatin et al., 2023; Zhou et al., 2023), the present work represents, to our knowledge, the first effort to integrate scRNA-seq data from neurodevelopmental-disease mice models.

When comparing ASD and control samples in our integrated dataset, we identified several differentially expressed genes (DEGs) across cell types at the third classification level. Notably, *Ttr*, which encodes transthyretin, was dysregulated in 13 clusters and emerged as the most upregulated gene in 7 of them, as shown in Figure 2A. *Ttr* encodes a non-canonical thyroid hormone transporter primarily responsible for carrying thyroxine (T4) across the blood-brain barrier (Richardson et al., 2015). Thyroid hormones are essential for brain development, neuronal differentiation, and synaptogenesis (Schroeder & Privalsky, 2014). Both maternal thyroid dysfunction during pregnancy (Kaplan et al., 2024) and individual thyroid dysfunction, particularly elevated T4 levels (Meng et al., 2024), have been linked to ASD. Beyond its role in thyroid hormone transport, *Ttr* is also involved in retinol (vitamin A) transport through its interaction with retinol-binding protein (Kang et al., 2018). Retinol is a crucial micronutrient for neurogenesis and neuronal plasticity (Shearer et al., 2012), and it has been implicated in ASD neurobiology (Liu et al., 2021), with emerging studies suggesting that vitamin A supplementation could serve as a potential therapeutic strategy for ASD (Guo et al., 2018).

Other consistently regulated genes include the murine pseudogene *BC023719*, a non-coding RNA downregulated in eight cell types. Although non-coding RNAs can regulate gene expression at both the transcriptional and post-transcriptional levels, there is limited evidence regarding *BC023719* function, apart from their expression in the central nervous system (CNS) and retina (Blackshaw et al., 2004). In contrast, members of the *Olig* gene family, which encode basic helix-loop-helix (bHLH) transcription factors, also showed consistent regulation across clusters. Notably, *Olig2* and *Olig1* were upregulated in five non-neuronal clusters (Mature Oligodendrocytes, Vascular Endothelial Cells, Pericytes, Oligodendrocyte Precursor Cells and Ependymal-Like Cells). *Olig* genes play a critical role in oligodendrocyte development and neural lineage specification, and changes in *Olig2* expression have been linked to ASD (Szu et al., 2021). Studies in VPA-induced ASD models have reported that *Olig2* expression in oligodendrocytes can be increased or decreased depending on the age (Bronzuoli et al., 2018; Graciarena et al., 2019), while a network analysis of gene expression of cerebellar tissue of ASD patients also shown up-regulation of *OLIG1* and *OLIG2* together with other oligodendrocyte markers (Zeidán-Chuliá et al., 2015), further supporting the relevance of these transcription factors in ASD pathology

An interesting observation from our analysis was the total number of DEGs per cluster, which revealed distinct transcriptional patterns among various cell types. Pericytes, astrocytes, and ependymal cells displayed a predominant downregulation pattern, whereas SST-like interneurons and OPCs demonstrated a strong upregulation trend. However, in most cell types, a balanced distribution of upregulated and downregulated genes was noted. Our analysis also identified a set of genes simultaneously enriched in the cell type and differentially expressed when comparing ASD vs control. For instance, in L4/5-IT excitatory neurons a significant positive correlation was observed. Those neurons play a crucial role in processing and relaying information between cortical areas (Im et al., 2022). The observed upregulation of genes that normally characterize these neurons suggests a potential intensification or dysregulation of their specific functions in the context of ASD. For example, the upregulation of genes encoding synaptic adhesion molecules (*Cntn6, Nrxn1, Cdh12*) and scaffolding proteins (*Dlg2*) could imply alterations in synaptic stability, specificity, or structure, potentially impacting cortico-cortical connectivity (Radies et al., 2012; Mercati et al., 2017; Cooper et al., 2024). Furthermore, the increased expression of genes modulating neuronal excitability, such as *Kcnip4* (Kv4 channel modulator) and *Hcn1* (HCN channel subunit), might directly impact the firing properties and integrative functions of L4/5 IT neurons (Marini et al., 2018; Ji et al., 2021). Such changes could contribute to broader cortical circuit dysfunctions and alterations in the excitation/inhibition balance, a frequently hypothesized mechanism in ASD.

In contrast, our analysis of astrocytes revealed a significant negative correlation: numerous genes highly enriched in astrocytes were significantly downregulated in ASD condition compared to controls. This widespread downregulation affects essential astrocyte markers and functional pillars such as *Aqp4* (aquaporin-4), *Kcnj10* (Kir4.1 potassium channel), *Gja1* (connexin 43), *Glul* (glutamine synthetase), *Atp1a2* (Na+/K+ ATPase alpha2 subunit), and the GABA transporter *Slc6a11*, strongly suggests compromised astrocyte function rather than hyperactivity in the ASD context. Aqp4 and Kcnj10 coordinately mediate water homeostasis and potassium buffering at the gliovascular interface and synapses; their downregulation likely impairs the clearance of extracellular K+, potentially contributing to neuronal hyperexcitability (Benga and Huber, 2012; Morin et al., 2020; Bonosi et al., 2023). Reduced *Gja1* expression indicates compromised astrocyte network communication through gap junctions, hindering spatial buffering and metabolic coupling essential for network stability (Altas et al., 2024). Similarly, the downregulation of *Glul* points to deficient glutamate-glutamine cycling, potentially leading to impaired clearance of synaptic glutamate (Fan et al., 2023), while reduced Atp1a2 suggests diminished capacity for the ion transport that powers neurotransmitter uptake (Sugimoto et al., 2020). This global downregulation of core astrocyte functional genes points towards a potential failure in glial support systems. Such astrocyte dysfunction is increasingly recognized as a contributing factor in ASD pathophysiology (Cano et al., 2024), likely to create a less supportive environment for neurons, exacerbating neuronal network instability, and contributing to the overall E/I imbalance observed in the disorder.

While the role of neurons in ASD etiology is well established, increasing evidence supports neuroinflammation and abnormal energy metabolism as contributing factors, indicating that non-neuronal cells such as astrocytes and oligodendrocytes play a crucial role in ASD neuropathology (Jiang et al., 2022). Particularly in mature oligodendrocytes, we found that 871 DEGs were found to be enriched in this cell type, and 91% of them were upregulated. When analyzing some of those genes using the IUPHAR and TF databases, several potential ASD-related targets emerged. For example, the upregulation of the prohormone convertase 1/3, encoded by *the Pcsk1 gene*, is a key enzyme involved in endocrine and metabolic regulation. Mutations in *Pcsk1* have been associated with gastrointestinal disorders (Stijnen et al., 2016), a condition frequently comorbid with ASD (Madra et al., 2021). Another important enzyme, peptidylprolyl isomerase A (*Ppia*), is linked to inflammatory diseases and has been identified as part of a gene set capable of distinguishing ASD patients from those with other developmental disorders (Hamzic et al., 2024). Additionally, the proto-oncogene *Jund*, a member of the JUN family, plays a crucial role in oxidative phosphorylation and has been reported to be upregulated in maternal immune activation mouse models of ASD (Li et al., 2022). This finding aligns with our GO enrichment analysis for mature oligodendrocyte DEGs, where “oxidative phosphorylation” (GO:0006119) emerged as the most significantly enriched biological process (BP). Oxidative phosphorylation is increasingly recognized as a critical pathway in ASD neurobiology, with evidence linking mitochondrial dysfunction to altered neuronal and glial energy metabolism in ASD (Carbonell et al., 2023). These results further reinforce the hypothesis that mature oligodendrocytes contribute to metabolic dysregulation and neuroinflammatory processes in ASD, enhancing our understanding of glial involvement in this pathophysiology.

While investigating the relationship among each reference dataset used in our integrated analysis, a strong correlation was observed between the subset of each reference and the integrated object for most cell types, reinforcing the validity of our integration approach and confirming that different ASD animal models, despite varying methods and paradigms, converge on similar molecular alterations. However, an intriguing exception emerged for vascular cells, which exhibited the lowest correlation and the fewest shared DEGs across models, except in the VPA model (Zhang et al., 2024). This suggests that vascular-related transcriptomic changes in ASD models may be model-specific rather than a broadly conserved feature of ASD pathology. While research on the role of vascular health in brain development and ASD is still emerging (Ouellette et al., 2024), previous studies indicate that VPA affects vascular development during early life (Manzo et al., 2025) and has anti-angiogenic effects in postnatal animals (Iizuka et al., 2018). Supporting this idea, subsets featuring specific neuronal mutations, such as Glutaminase 1 deficiency in *CamkIIa+* cells from Ji et al., 2023, did not exhibit significant vascular alterations, further reinforcing that vascular dysregulation may be linked to specific environmental or pharmacological models.

Furthermore, comparing DEGs from our integrated dataset with those reported by Donnard et al., 2022 in a Fragile X Syndrome (FXS) model revealed substantial similarities in gene regulation, suggesting overlapping molecular pathophysiology between FXS and the diverse ASD models integrated here. In the original work from Donnard et al., 2022, they notably reported an astrocyte-mediated exacerbation of excitatory-inhibitory imbalance. Aligning with this, astrocytes showed the highest number of shared DEGs when comparing their dataset with our integrated results. This concordance extended to correlation analyses performed between data subsets (from our integration *vs*. Donnard et al., 2022 data subset), where neuronal and astrocyte populations exhibited the strongest correlations, while immune cells displayed lower transcriptional similarity. Moreover, as we previously reported, 98% of DEGs common with enriched genes in astrocytes were found to be downregulated. This pronounced downregulation points towards significant deficits in astrocyte function within these models, reinforcing the crucial role astrocytes may play in ASD-related circuit dysregulation, as highlighted by others (Talvio and Castrén, 2024).

As previously mentioned, the imbalance between excitatory and inhibitory signaling is a leading hypothesis in ASD neurobiology (Uzunova et al., 2016), and this was strongly reflected in our cell-cell communication analysis, which revealed increased neuronal communication in both excitatory and inhibitory populations. This pattern was also observed in the specific neuronal communication analysis of human single-cell data (Zhao et al., 2023), further supporting our findings. Notably, NicheNet analysis identified several ligands and predicted target genes that align with pathways exhibiting increased communication in ASD, as revealed by CellChat analysis, particularly PTN, SLIT, and PTPR signaling. Among the top-ranked ligands, *Pdgfa* showed significant activity and upregulation in ASD, consistent with the role of the PTN pathway in growth factor signaling and extracellular matrix (ECM) regulation (González-Castillo et al., 2015). Interestingly, PTN has been demonstrated to bind to VEGFR2, thereby inhibiting VEGFA signaling (Lamprou et al., 2020), one of the predicted ligands identified in our ligand-receptor analysis. Notably, previous research found that blocking VEGFA alleviated non-vascular FXS abnormalities, such as cognitive impairments (Belagodu et al., 2017). Moreover, predicted target genes such as *Ncam1* and *Col4a1* are also involved in ECM remodeling and cell adhesion (Kuo et al., 2012; Vukojević et al., 2020), further reinforcing the significance of these processes in neuronal dysregulation found in ASD.

In addition to PTN signaling, *Reln* (Reelin), another predicted ligand, plays a critical role in axon guidance and synapse organization (Faini et al., 2021), suggesting a potential link to the SLIT pathway, which is essential for neuronal wiring and connectivity (Gonda et al., 2020). Interestingly, both *Reln* and *Robo1* (a key gene from the SLIT/ROBO pathway) have been shown to modulate neuronal adhesion via N-cadherin interactions (Rhee et al., 2007; Matsunaga et al., 2017), further supporting their involvement in neurite outgrowth and synaptic organization. Moreover, *Adamts3*, a metalloproteinase identified as one of the top ligands in our analysis, is known to inactivate Reelin (Ogino et al., 2017) and has been previously linked to ASD through GWAS studies (Rexrode et al., 2024), reinforcing the idea that dysregulation of *Reln* processing may contribute to altered neuronal development in ASD (Scala et al., 2022). While direct components of the SEMA3 signaling pathway were not identified, a closely related semaphorin family member, *Sema4g*, was found in the ligand-receptor analysis. Semaphorins, in general, have been consistently associated with ASD (Steele et al., 2021; Carulli et al., 2021), and although the specific function of *Sema4g* remains unclear, it may play a role in axon guidance (Belyk et al., 2015).

Remarkably, a key receptor in the PTPR signaling pathway, *Ptprz1*, was found to be downregulated in L5/6 NP and other excitatory neurons, while its predicted ligand, *Cntn1*, was downregulated in other inhibitory neurons (Supplementary Table 6). Although *Ptprz1* and *Cntn1* are well known for their roles in neurite outgrowth and cell adhesion in glial cells (Mohebiany et al., 2012), their involvement in neuronal signaling remains largely unexplored. However, a recent study identified *Cntn2* as an interaction partner in a gene-gene *in silico* analysis for both *Cntn1* and *Ptprz1*, suggesting that these molecules form a functional signaling complex in patients with idiopathic generalized epilepsy (Lin et al., 2024). Given that epilepsy is a common comorbidity in ASD (Keller et al., 2017), this finding indicates that PTPR signaling dysregulation may contribute to both ASD and epilepsy-related neural dysfunction.

As anticipated, a significant overlap was found between the DEGs identified and the genes cataloged in the SFARI database for both human and animal models. A key advantage of our study is that it allows us to explore these genes at a single-cell level. For example, *Ermn*, which encodes the oligodendroglia-specific cytoskeletal protein Ermin, was upregulated across all non-neuronal populations. Rare genetic variants leading to hypomethylation at the *ERMN* locus have been significantly associated with ASD patients (Homs et al., 2016), which corroborates with our findings of *Ermn* upregulation in non-neuronal cells. However, while *ERMN* expression was found to be reduced in peripheral blood cells of an ethnic cohort of ASD patients (Shiva et al., 2021), this discrepancy may be due to differences between peripheral and central nervous system expression patterns.

Another key gene, *ATP2B2*, encodes calcium-transporting ATPase-2, an essential Ca²⁺ extrusion pump in neurons. Variants in *ATP2B2* have been associated with ASD-like phenotypes in transgenic mice, which present calcium extrusion dysfunction, motor impairments, and cerebellar atrophy (Poggio et al., 2023). Furthermore, *ATP2B2* loci polymorphisms have been consistently linked to ASD in large-scale GWAS studies (Autism Spectrum Disorders Working Group of The Psychiatric Genomics Consortium, 2017). Notably, in our dataset, *Atp2b2* was consistently downregulated across multiple neuronal subtypes (L2/3 IT, L4/5 IT, L5 IT, Other Excitatory Neurons, Other Inhibitory Neurons, SST-like). Given its vital role in calcium homeostasis, this downregulation is likely to directly influence neuronal function, potentially contributing to ASD-associated endophenotypes.

Other key genes identified in our analysis that may be associated with specific neuronal subtypes include the transcription factors (TFs) *FOXG1* and *MEF2C*. Mutations in *FOXG1* are strongly linked to FOXG1 syndrome (Wong et al., 2019) and have also been implicated in Rett syndrome (RTT) (Mazel et al., 2024), both of which are characterized by severe developmental delays and cognitive impairments. In our dataset, *Foxg1* was found to be downregulated in L5/6 NP neurons, a population that comprises the deep layers of the cortex. *FOXG1* belongs to the forkhead transcription factor family and is one of the earliest TFs induced in the neural progenitors of the forebrain cells. It plays a crucial role in post-mitotic neurons, particularly in cortical laminar organization (Hettige and Ernst, 2019). Consistently, it was demonstrated that *Foxg1* in pyramidal neurons is essential for establishing cortical layers and the identity/axon trajectory of callosal projection neurons. This occurs through the formation of a complex with *Rp58* that directly represses genes such as *Robo1*, *Slit3*, and *Reelin*, regulators of neuronal migration and callosal axon guidance and relevant targets found in our communications analysis. Importantly, the inactivation of just one *Foxg1* allele specifically in cortical neurons is sufficient to cause cortical hypoplasia and corpus callosum agenesis (Cargnin et al., 2018), explaining core features of FOXG1 syndrome and highlighting the relevance of its downregulation observed in our data.

Similarly, *MEF2C* is a TF frequently associated with ASD and monogenic disorders that mimic RTT, playing a key role in neurogenesis and synaptic pruning (Zhang & Zhao, 2022). In our dataset, *Mef2c* was found to be downregulated in MEIS2-like interneurons, suggesting a potential functional link between these transcription factors in neuronal differentiation and synaptic remodeling. Interestingly, mutations in *MEIS2*, another TF, have also been linked to clinical features overlapping with RTT (Srivastava et al., 2018). Further emphasizing the critical neurodevelopmental roles of these specific TFs, studies utilizing whole-genome sequencing to precisely map breakpoints in patients with balanced chromosomal rearrangements and associated phenotypes like intellectual disability have directly implicated both *MEF2C* and *MEIS2*. These analyses established diagnoses by identifying cases where *MEF2C* or *MEIS2* were directly disrupted by the chromosomal breakpoint (Schluth-Bolard et al., 2019). These findings highlight how TF-mediated regulation may contribute to altered cortical organization and neurodevelopmental deficits observed in ASD pathophysiology, particularly in neuronal subtype-specific contexts.

Finally, our comparative analysis of the integrated object and human postmortem datasets across various cortical regions provides strong evidence of conserved transcriptomic alterations in ASD while highlighting region- and cell-type-specific differences. A substantial overlap of DEGs was found, particularly in the prefrontal and parietal cortices, which have been widely implicated in ASD (Wymbs et al., 2021; Leisman et al., 2023). The parietal cortex exhibited the highest degree of consistency, with notably conserved and positively correlated transcriptional regulation across excitatory neurons (874/1198 shared DEGs), inhibitory neurons (196/285 shared DEGs), and mature oligodendrocytes (496/627 shared DEGs).

SynGO analysis revealed that in both excitatory and inhibitory neurons, pathways related to synaptic signaling and protein synthesis were significantly enriched, particularly “translation at presynapse” and “translation at postsynapse”. This highlights potential vulnerabilities in synaptic function fundamental to neuronal communication. For instance, altered expression of presynaptic genes like *CACNA1A*, crucial for the release-triggering Ca2+ influx in excitatory neurons whose mutations are linked to neurodevelopmental disorders (Kramer et al., 2023), or *ATP6V0A1*, essential for neurotransmitter loading into vesicles in inhibitory neurons, vital for GABAergic function and often implicated in epileptic encephalopathies (Bott et al., 2021), could directly impair neurotransmission and disrupt the critical excitation/inhibition balance. Similarly, dysregulation of postsynaptic components, such as the NMDA receptor subunit *GRIN2B*, a well-established ASD risk gene (Pan et al., 2015) or the AMPA receptor regulator *CACNG2* in excitatory neurons, points towards disrupted signal reception and plasticity mechanisms (Lee et al., 2025). In inhibitory neurons, alterations in genes like the signaling molecule NRG1, known to regulate GABAergic circuit development and function, further underscore potential disruptions in inhibitory control (Navarro-Gonzalez et al., 2021). The consistent finding of CAMK2A alterations, a key kinase involved in synaptic plasticity, across both neuron types in postsynaptic analyses suggests a broadly impactful disruption of signaling pathways critical for learning and memory, often affected in ASD (Yassuda et al., 2022).

The enrichment in presynaptic active zones and postsynaptic density components further supports the hypothesis that abnormal synaptic architecture contributes to the pathophysiology of ASD. Crucially, the identification of ribosome-related components and the explicit enrichment of pathways like “translation at presynapse” strongly corroborated by the altered expression of numerous ribosomal protein genes (e.g., *RPL* and *RPS* genes) found specifically in the inhibitory presynaptic dataset of this study aligns robustly with previous findings indicating widespread alterations in ribosomal gene expression and protein synthesis machinery in both postmortem cortical tissue and iPSC-derived cells from ASD patients (Lombardo et al., 2021). This convergence suggests that disruptions in synaptic protein homeostasis, potentially affecting both global translation and the critical local translation required for synaptic maintenance and plasticity, may represent a conserved molecular feature contributing to ASD pathophysiology (Porokhovnik et al., 2015).

For mature oligodendrocytes, Metascape analysis using the DisGeNET platform demonstrated a significant enrichment of disease-associated genes, particularly those connected to neurodevelopmental disorders, epilepsy, and mental disorders. Considering the role of oligodendrocytes in axonal myelination and neuronal connectivity, disruptions in oligodendrocyte maturation and functional processes could lead to deficits in neuronal circuit stability and information processing in ASD (Gálvez-Contreras et al., 2020). Moreover, the involvement of NRXN and DLGAP family genes in our analysis indicates that oligodendrocytes may directly influence synapse stabilization and plasticity, reinforcing previous findings that white matter abnormalities in ASD are associated with altered NRXN signaling in oligodendrocyte function, demonstrating that by saturating NRNX with exogenous soluble neuroligin protein blocks axo-glial signaling by oligodendrocytes and axons in both *in vitro* and *ex vivo* assays (Proctor et al., 2015). These results further support the increasing recognition that glial dysfunction plays a crucial role in ASD beyond its traditional function.

Our integrated approach enabled a robust identification of conserved DEGs across diverse models, as illustrated in Figure 2, while also allowing direct comparisons of gene expression patterns among models with distinct experimental paradigms (Figure 3). Despite the strengths of this study, several limitations must be acknowledged. First, we cannot completely exclude the possibility that integrating scRNA-seq datasets from different ASD genetic and pharmacological models introduces variability due to differences in experimental protocols, developmental stages, and brain regions. While bioinformatics approaches were employed to mitigate batch effects, residual variability may still influence the findings (Luecken et al., 2022). Second, ASD is a highly heterogeneous disorder, and although this study identifies conserved transcriptional alterations, it may not capture all aspects of its complexity (Martinez-Murcia et al., 2017). Moreover, more data is available for other animal models built on ASD risk genes but we opted to focus on specific models here. Additionally, comparisons with human postmortem data are limited by differences in developmental timing and confounding factors like chronic clinical treatment (Fetit et al., 2021). Future studies should explore longitudinal analyses and functional validation to elucidate further the causal relationships between the identified transcriptomic alterations and ASD pathogenesis.

In summary, our findings of increased interactions between inhibitory and excitatory neurons suggest widespread network-level dysfunction, reinforcing the hypothesis of an altered excitatory-inhibitory balance in ASD. Moreover, non-neuronal cells like astrocytes and mature oligodendrocytes could play a major role in this dysfunction, as evidenced here and suggested by others. Comparative analyses with human postmortem datasets further validate these findings, demonstrating the translational relevance of our integrated approach. Only through this integrated approach is it possible to simultaneously increase the statistical power and relevance of our findings, as well as enable comparative analyses among different models. We believe that DEGs consistently regulated across different experimental paradigms in animal models, and which show similar patterns in corresponding cell types in ASD patient samples, are pivotal to cellular brain function and essential for understanding the neurobiology of ASD.

