## Supplemental Figure for "Integrative Single-Cell Analysis of Autism Spectrum Disorder Animal Models Reveal Convergent Transcriptomic Dysregulation Involved in Excitatory-Inhibitory Imbalance and Glial Disfunction"

*Corresponding author:

SUPPLEMENTARY FIGURES

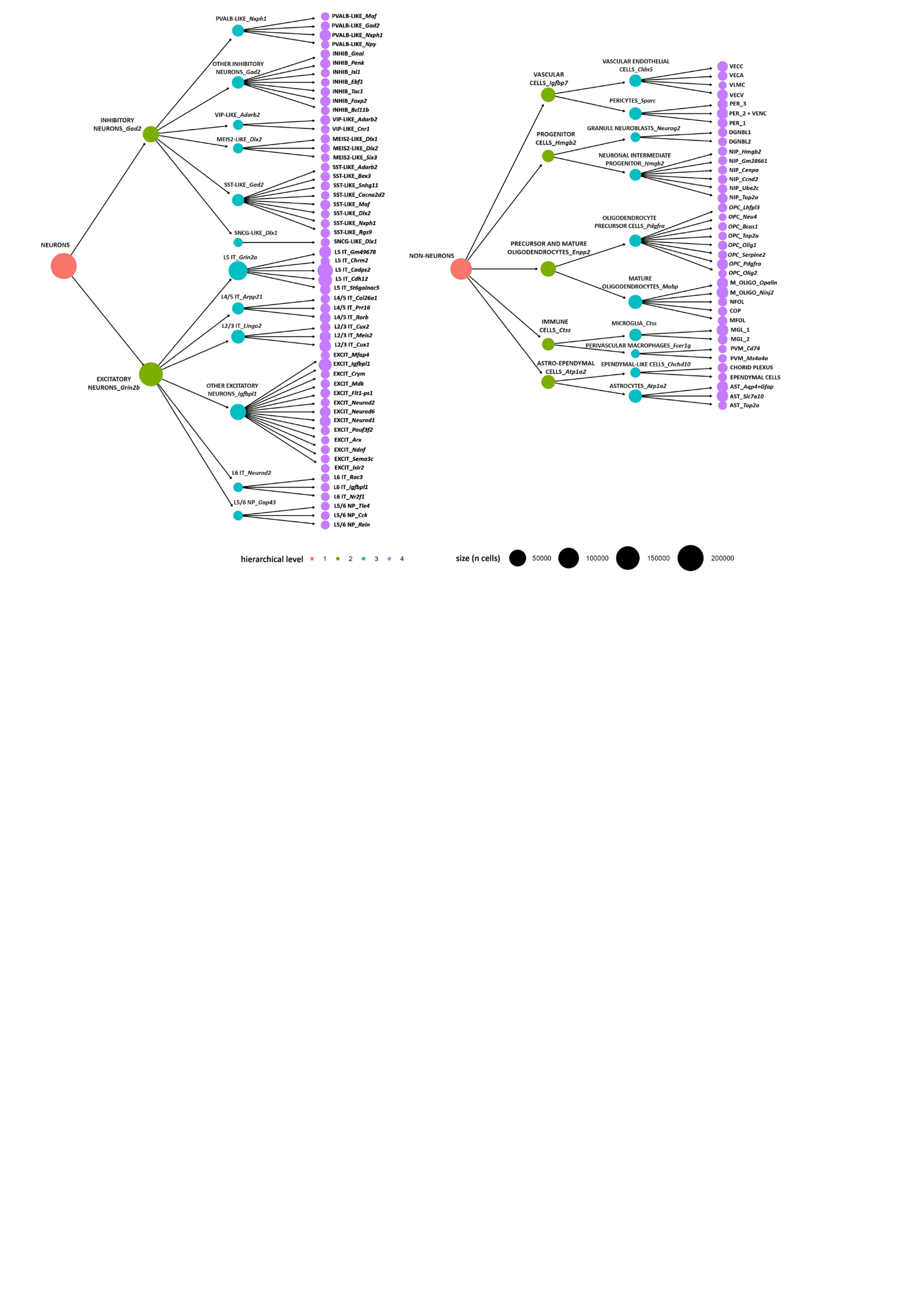

**Supplementary Figure 1**. Hierarchical dendrogram illustrating the clusterization process up to the fourth level. Levels 1, 2, 3, and 4 are shown, with the second and third levels annotated with the most differentially expressed gene for each cluster, while in the fourth level we maintained a functional nomenclature or added the most differentially expressed gene in that cluster.

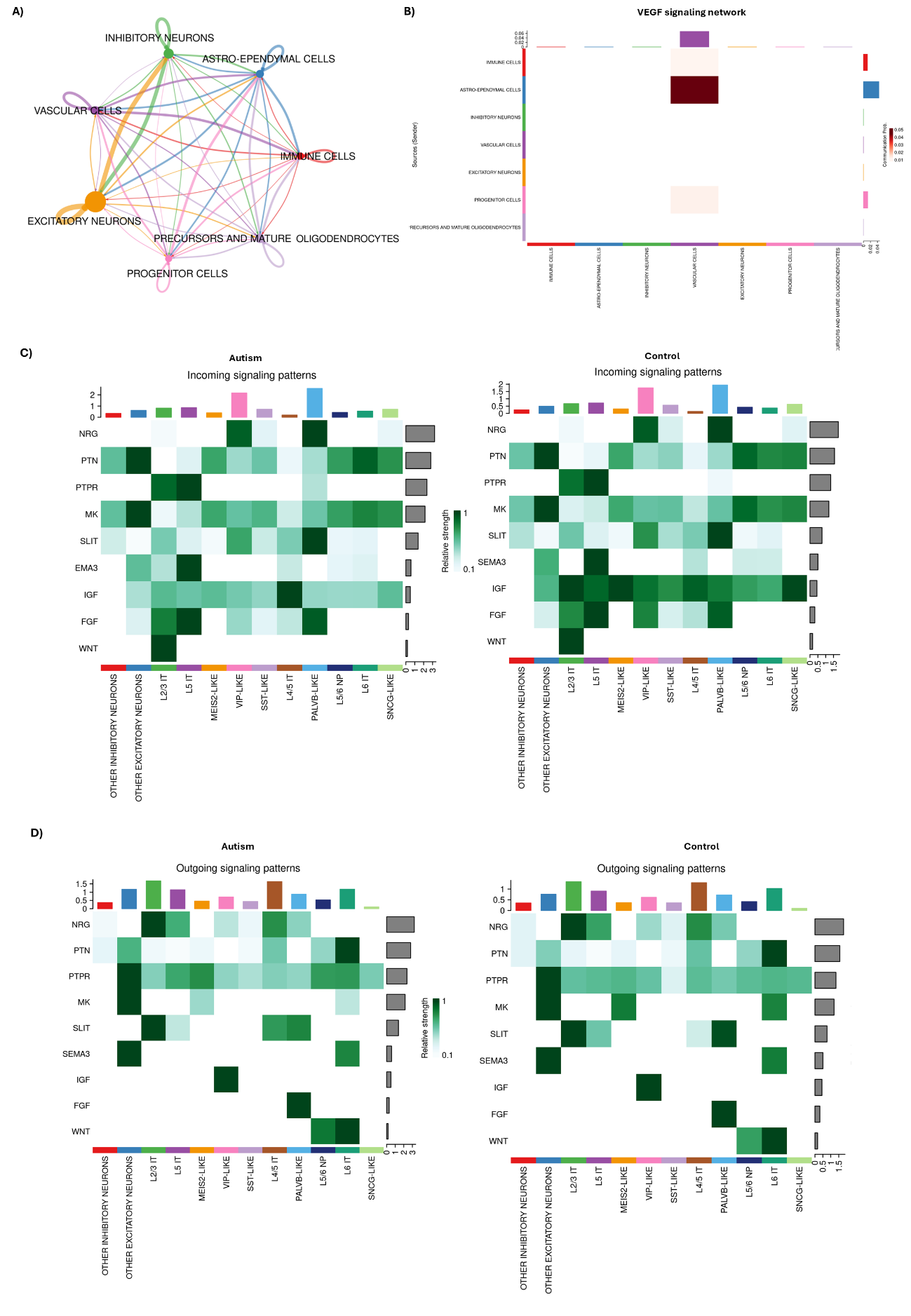

**Supplementary Figure 2**. **CellChat Analysis of Cellular Communication in the Integrated Dataset and Condition-Specific Comparisons (Autism vs. Control). A)** Overall cellular communication network for all cell types in the integrated dataset, independent of autism or control conditions. Nodes represent cell types, and edge thickness reflects the strength of signaling interactions. **B)** VEGF signaling network showing the key sender and receiver cell types. The heatmap highlights the contributions of vascular cells and astro-ependymal cells as primary participants in VEGF signaling pathways. **C)** Heatmaps showing incoming signaling patterns (receiver signals) across neuronal subtypes in autism (left) and control (right). Differences in pathway activity highlight the dysregulation of specific signaling pathways in autism. **D)** Heatmaps showing outgoing signaling patterns (sender signals)

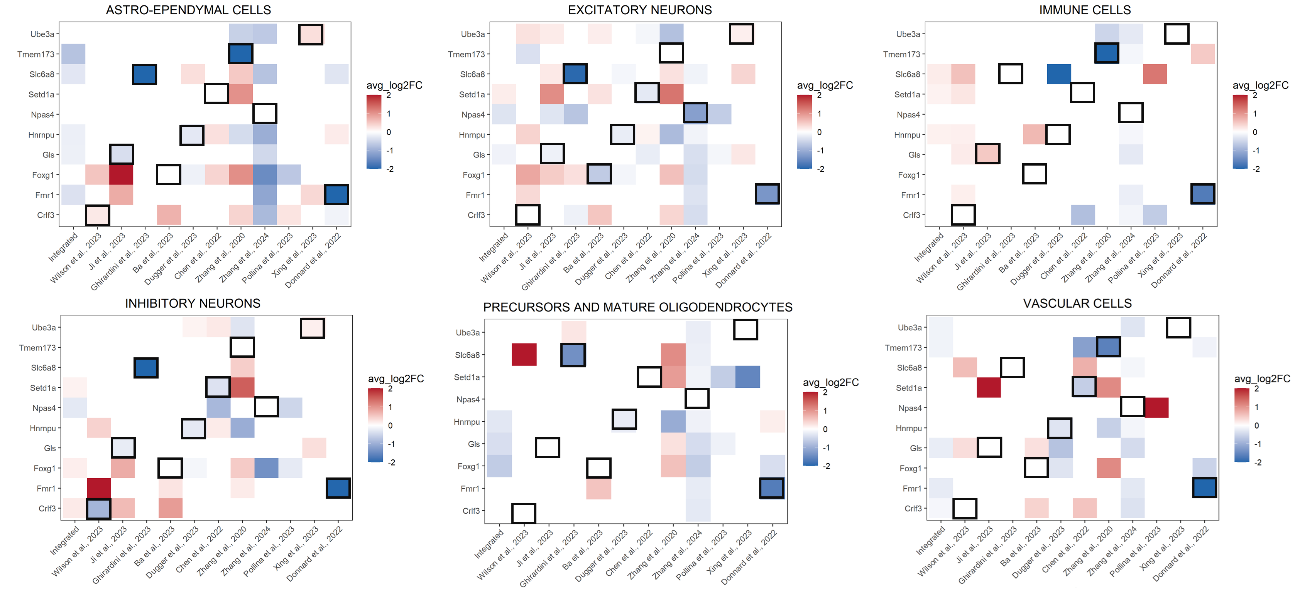

**Supplementary Figure 3. Heatmaps depicting differentially expressed genes in the integrated object and the subset of each study across distinct cell types:** Each column represents a different reference (study), while each row corresponds to a gene of interest (modulated in each model). The color scale reflects the avg_log2FC values, ranging from blue (negative) to red (positive), with white cells indicating non-significant or absent data for that gene in each reference. Each highlighted square indicates the correspondence of the modulated gene to the reference in that column.
